# Clustered Loss of Dendritic Spines Characterizes Encoding of Related Memory

**DOI:** 10.1101/2020.12.17.423264

**Authors:** Suraj Kumar, Meenakshi Prabod Kumar, Yagika Kaushik, Balaji Jayaprakash

**Affiliations:** Centre for Neurosciences, Indian Institute of Science, Bangalore – 560012.

## Abstract

Generation of new spines is often thought of as a correlate of memory and loss of spines is considered representing memory loss. Contrary to common belief, we observe that spine loss has functional value in distinctly encoding related life events rather than causing memory loss. Using spatial autocorrelation of dendritic morphology obtained from in vivo longitudinal imaging, we show that clustered loss, rather than gain, of new spines characterizes the formation of related memory. This spatially selective dendritic spine loss occurs closer to new spines formed during the acquisition of initial memory. Thus, enabling the dendrites to store multiple memories and their inter relationship. Remarkably, we find acquisition of related memory in the absence of NMDAR activation increases the fraction of such correlated spine loss.

## Main Text

Neuronal plasticity is known to span several scales and elucidating its physical substrates has been an intensely researched area of neuroscience. Cajal^1^ postulated dendritic spines, protrusions in dendrites that represent putative synapses in an excitatory neuron, as physical substrates of memory. Since then, several studies have shown that spine density has been positively correlated with memory, while spine loss is associated with LTD and memory loss^2–9^. Further, studies on artificial intelligence have negated pruning based learning models outside of development as they result in catastrophic forgetting^10,11^. Advent of two-photon microscopy enabled studying these spines in live brain slices following long-term potentiation (LTP), and a correlated increase in spine density was seen^12–15^. Longitudinal in vivo imaging^16–19^ established that spines are dynamic, and long-term stability amongst them could represent plastic events. Similarly, spine densities have been shown to be correlated with motor learning, and a different set of spines were seen representing different motor skills^9^. Thus, prevailing evidence strongly suggests memory formation is accompanied by new spine generation with no clear role identified for the spine loss.

Memories of life events are acquired, stored, and consolidated through a cascade of processes engaging molecules, cellular ensembles, and circuitry of the network^20^. Recently it has been shown that temporally connected memories are stored in overlapping but different ensembles of neurons^21^. However, memories that are related through their contents have been shown to be encoded independent of NMDA receptor activation but only if a prior related memory is already acquired^22–24^, suggesting that the neurons encoding a prior memory could possibly act as a substrate for new but related memory. In this context, it is of interest to know how memories that are related in their content are encoded, preserving their relation and their individual identity simultaneously. Dendritic spines represent an intriguing possibility for encoding such memories. Studies involving cortical areas that control fear expression and extinction show changes in spine density following memory formation^25,26^. A decrease in spine density in the CA1 region of the brain has been reported^27^ in conjunction with memory formation. However, it is unclear if spine loss occurs in a functionally relevant manner, especially when a new memory formation is positively correlated with spine genesis. Further, the spatial organization of spines has been shown to provide additional dimensions for the neurons to enhance its storage capacity^28–30-^. Such clusters have been shown to play a vital role in dendritic computation, and recent findings have shown the occurrence of dendritic action potentials from human brain tissue^30^. However, it is not known if the clustered plasticity can be a substrate for related memory, and if spine loss has a functional role in encoding memory.

We develop spatial auto-correlation function (ACF) as an analytical tool for tracking the morphological changes that occur along the length of the dendrites. Through longitudinal *in vivo* imaging, contextual fear conditioning, and ACF, we questioned if information could be encoded in the spatial organization of spines. We studied the changes that accompany the formation of new memory as well as related memory in the presence and absence of NMDAR blockade. We find that mice, when trained in behavioral scheme (Fig. 1A) involving one-shot learning, CPP effectively blocks learning, and this blockade can be rescued if the animals had prior training in a related but different context. Freezing responses (Fig. 1D) obtained from the animals trained in first context (Ctxt A) in presence (Grp 2) and absence of CPP (Grp 1) showed that the learning is blocked in Grp 2 as expected^22–24^. The same set of animals when trained in second context (Ctxt B, related memory) but now with the Grp 1 receiving the CPP while Grp 2 receiving saline exhibited freezing for the training context in both groups. We interpret this as the rescue of CPP induced acquisition blockade by prior learning. We note that in our design the groups act as their internal control for the efficacy of the blockade. Consistent with previous^22–24^ studies we see that memory of second context is acquired by both the groups. We argue in such a design, acquisition of context A would reflect new memory and context B would correspond to related memory as it is acquired in the absence of NMDAR activation but only when memory for similar events is present initially. It is important to note that the memories acquired for both the contexts have shown to be distinct previously^23^. Next, we focussed on following the spine level changes that accompany such learning events.

**Figure 1:**
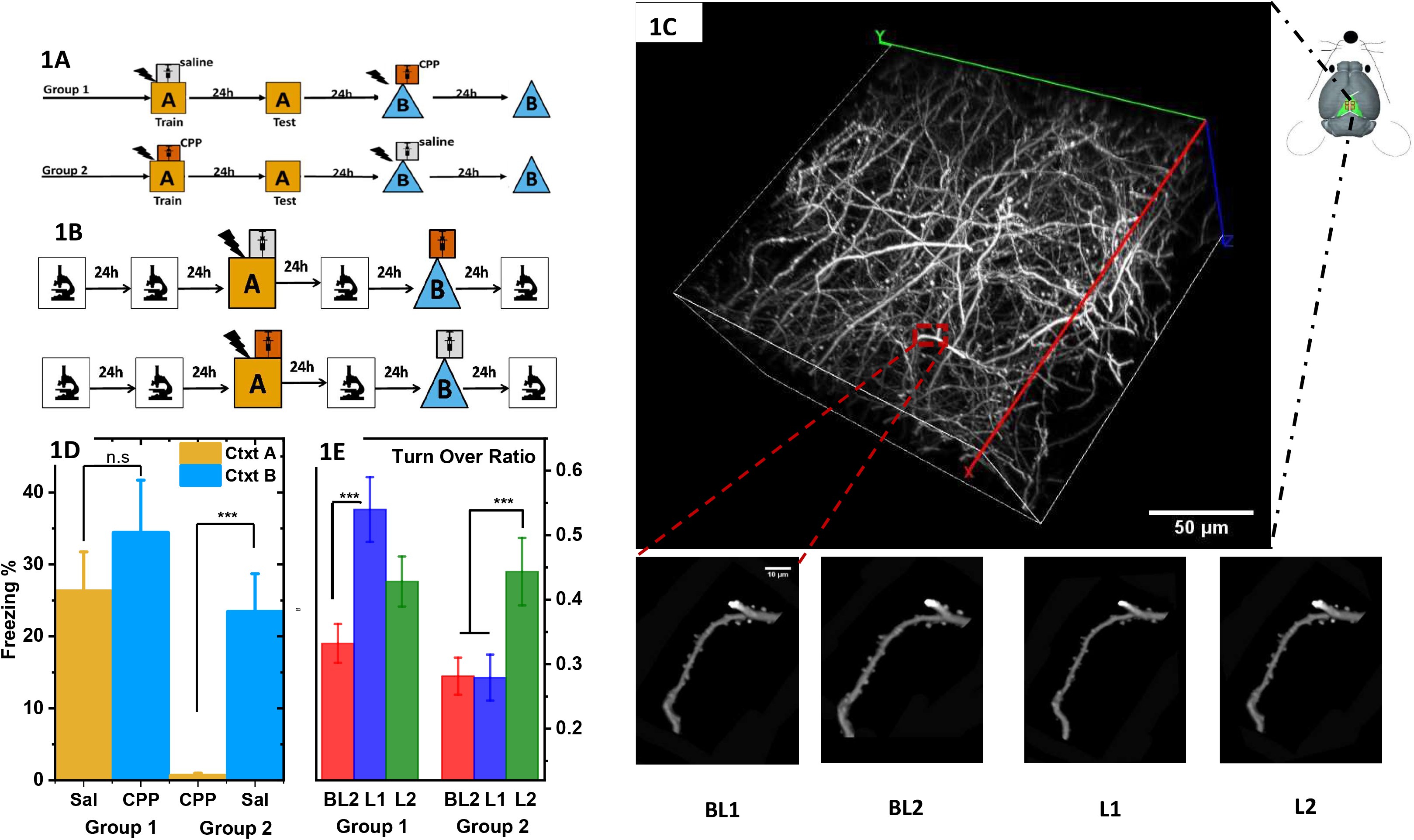
Learning of novel context and not a related context causes an increased spine turnover. (A) Behavioral scheme used for training the mice in novel and related contexts. Two groups of mice are trained in context A one administered with saline and other with CPP. Same set of mice underwent training in context B but now the animals that received saline previously are administered CPP while that received CPP is administered saline. (B) Schematic representation of the imaging sessions and the timeline of training. Note that the mice that underwent imaging are not used for retrieval to exclude retrieval induced changes. (C) Mice brain (200 × 200 × 150 μm) imaged in vivo is reconstructed in 3D to show the neuronal architecture. The scale bar is 50 μm. Location of the imaging area (RSc) in the mice brain is shown as an illustration. The area shown within red square is enlarged to show the spines located on the dendrites in images obtained at 4 time points (Baseline1(BL1), Baseline2(BL2), First-Training(L1) and Second training (L2). Scale bar in the enlarged image corresponds to 10 μm. (D) Comparison of freezing exhibited by mice from both these groups show that difference among the groups is significant (ANOVA, F> 7, p < 0.001). Post-hoc analysis shows that mice (n = 9(group1), 8(group2)) freeze significantly less when CPP is administered (0.64 ± 0.35) compared to saline (26.5 ± 5.5) in context A (p< 0.01). These mice show comparable freezing (p> 0.46) when subsequently trained in context B for both saline (23.4 ± 5.3) and CPP (34.4 ± 7.3) administration. Thus, CPP is effective in blocking the acquisition of contextual memory and prior learning rescues such deficit. (E)Spine turnover, defined as fraction of spines undergoing a change, is significantly different across the sessions in both these groups (ANOVA: Group1(N = 103) ,F>6.5, p < 0.0016, Group2(N = 70), F> 5.4, p < 0.0048) post hoc analysis showed that it is significantly higher at L1 in group1(Bonferroni, p< 0.001 (L1), p<0.216(L2)) shown as blue 0.54 ± 0.05, and not L2 (olive bars, 0.43 ± 0.04 in left). However, in group 2, only L2 (shown as olive bar (0.44 ± 0.05)) and not L1 (blue bar 0.28 ± 0.03) is higher (Bonferroni, p < 1(L1), p< 0.014(L2)). as compared to the baseline (red bar (group 1, 0.33 ± 0.03) (group 2, 0.28 ± 0.029)).

Immediate early genes-based studies have established the role of retrosplenial cortex (RSc) in encoding and storage of contextual memory^31,32^. Schema formation is thought to be consolidated representation of several related information involving RSc activity^33^. In RSc, acquisition of contextual memory has been shown to simultaneously produce clustered addition of new spines^34^. Hence, we used our behavioral design interspersed with *in vivo* imaging of RSC (Fig. 1B) to follow the dendritic changes accompanying training in new and related contexts. Our imaging scheme (Fig. 1B) ensures the observations can be attributed to training and not to retrieval. Images of Thy-1 YFP mice from four imaging sessions (~300 dendrites from 12 ROIs and 6 animals across both the groups) are aligned and same dendrites are identified in each session. Figure 1C shows the reconstructed brain volume and panel below shows zoomed region of one dendrite extracted from longitudinal images. The mice are imaged four times, twice before training (BL1, BL2), once after training in first context(L1) and again following second context training (L2). We characterize the spine dynamics in these two groups and find the turnover increases following learning compared to baseline (Fig. 1E). Spine level changes, turnover and not density (SFig. 1) is seen following new learning.

Previously dendritic spines have been shown to cluster following formation of new memory^9,34,35^. Identification of such spine clusters is often confounded by changes in spine density and limited sensitivity of measuring changes in thresholded cumulative frequency distributions (CFDs). While these methods serve to identify clusters in an efficient manner, the use of a single distance for a given spine is restrictive and prevents further characterisation.

Auto-correlation functions (ACF) are an effective way of characterising random phenomena. We argue that if clustering of spines were to occur, a more sensitive way to detect clustering would be to compare interspine distances over the entire dendrite. We constructed a digital sequence corresponding to each dendrite as explained in the methods (SI). Briefly, the dendrite is discretized into small units of uniform length and the presence or absence of spine in this unit is indicated by “1” or “0” respectively (Fig. 2A). Length of the unit is such that it contains utmost one spine and a given spine is contained within a fragment. These sequences uniquely represent each dendrite and are used for calculating the ACF. Since autocorrelation measures the similarity of given point in a data-train across different delays, we argue that clustering would reflect as prolonged decay of the correlation. To quantify the extent of clustering from the observed data we develop an analytical description of these sequences (SI:Theory). We model the spatial arrangement of spines as originating from an underlying Poisson process and reason that having found a spine at start position(l=0), the probability of finding another spine at “l > 0” given a mean spine density of “μ” is equivalent to asking how long one needs to wait for arrival/occurrence of an event in a Poisson process with a mean rate of “μ”. Such wait times are Erlang distributed and is given by,

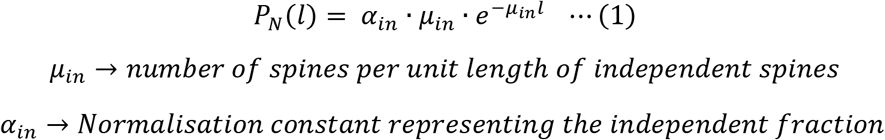

**Figure 2:**
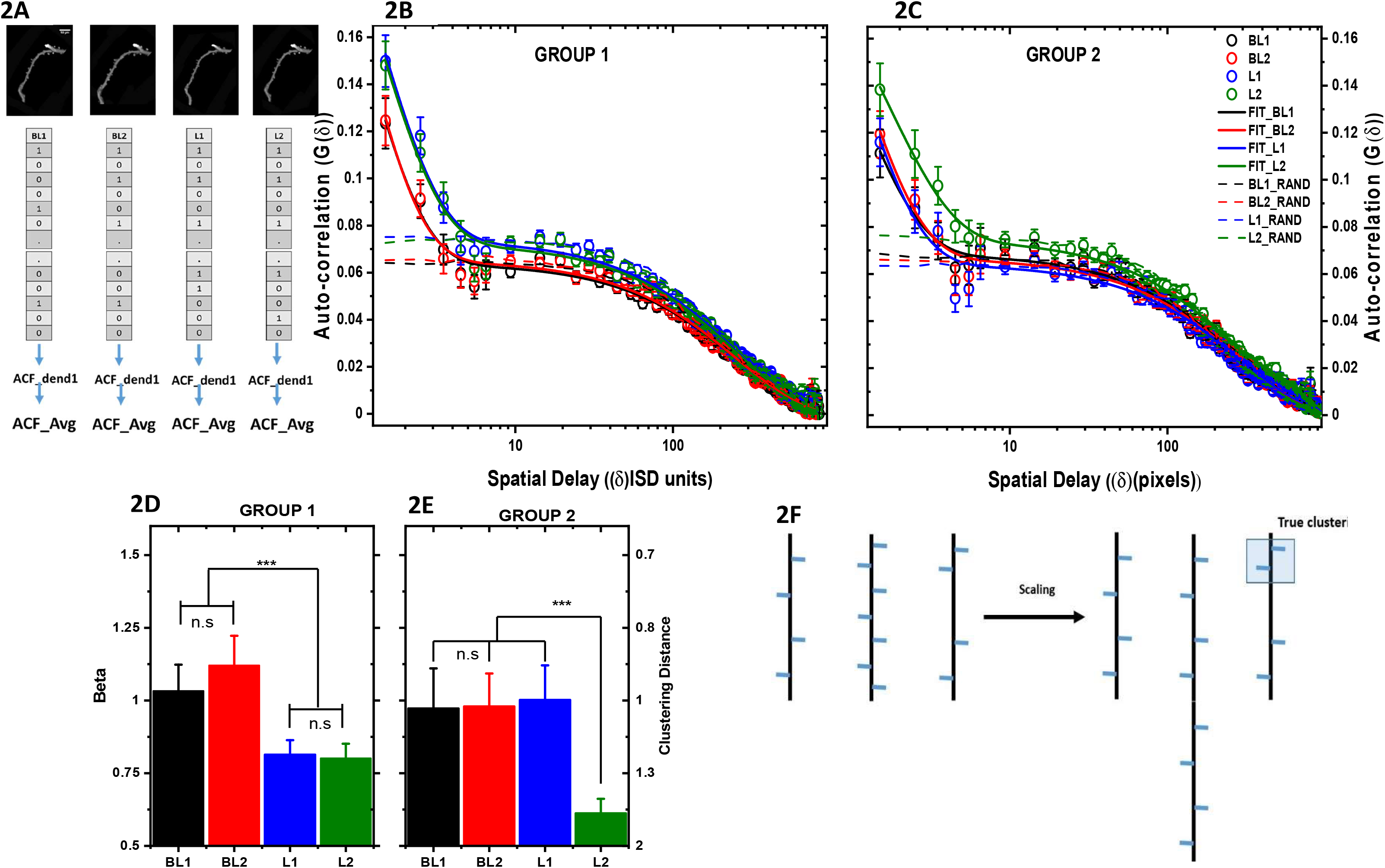
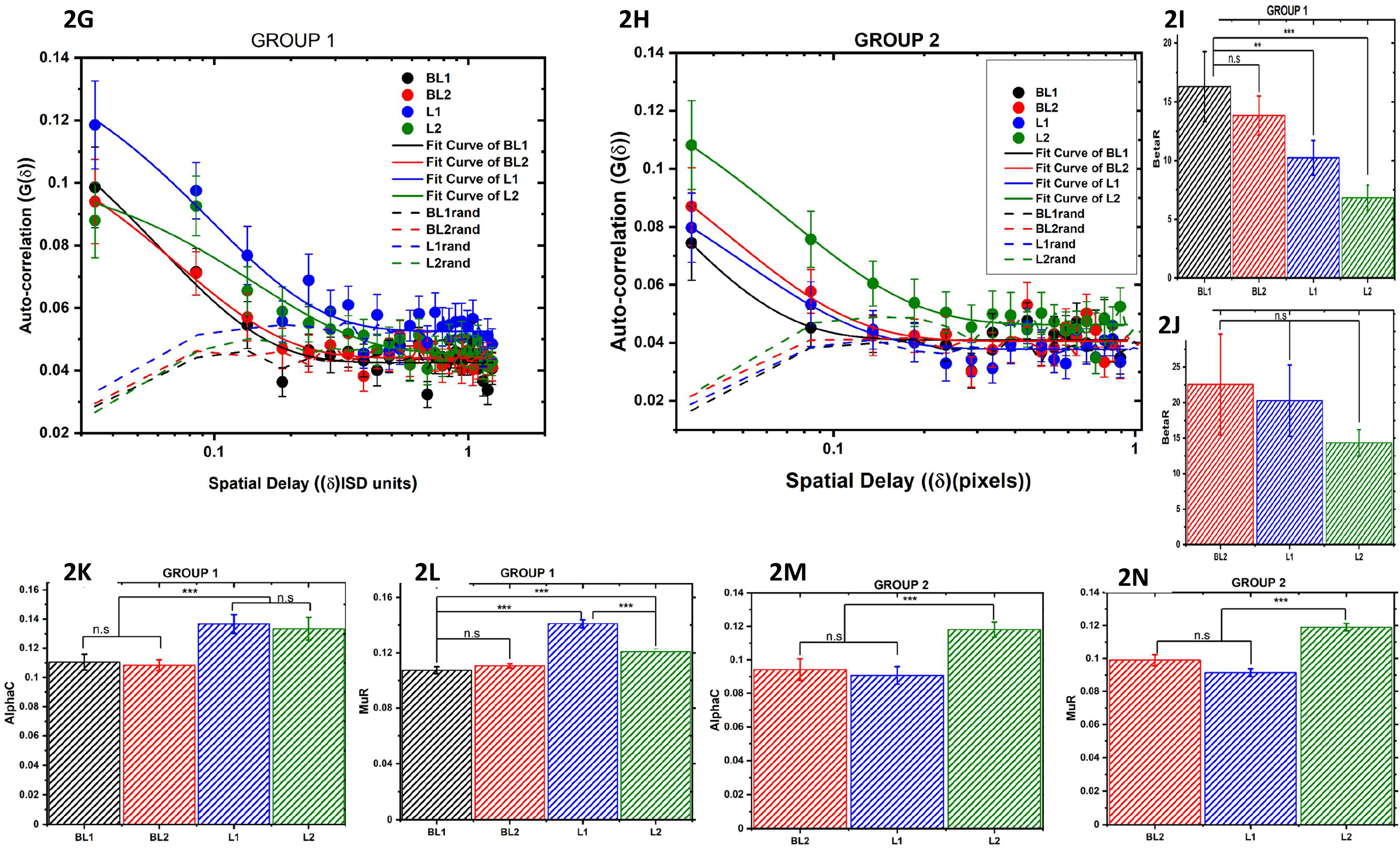
Related learning decreases the normalised clustering distance without altering the clustering fraction. (A) Schematic on ACF generation workflow. The images of dendrites are discretised such that a unit utmost can have one spine and any given spine is contained in just one unit. These sequences of units are then used for measuring the ACF and the resulting ACF is averaged across dendrites. (B)The open circles are the spatial ACF generated from 103 dendrites during sessions BL1(black), BL2 (red), L1(blue) and L2(olive). This colour scheme is maintained through the text. The ACFs differ significantly at shorter distances (τ < 125) when tested with non-parametric Friedman ANOVA (Chi-Sq = 73, DF = 3, p<10^−16^). Post-hoc analysis (Table 1A) revealed that both L1 and L2 are significantly different than BL1 /BL2(Z > 5, p< 10^−6^). The dashed lines are the session wise ACFs obtained after shuffling the location of spines along a dendrite. Pair wise comparison of the dendritic ACF and their corresponding shuffled sequence is found to be significantly different when compared with Friedman’s ANOVA. The pairwise statistics with its p values are listed in Table-1B of SI. L2-L2Shuffled comparison has the highest chance factor (p < 0.01) among the four. Solid lines are the fit of the session-wise ACF to Eq. 3. The data fit well to the above equation (Adj. R. Sq > 0.9). The fit parameter β across sessions are compared in (D) and (E) for group1 and group2. The difference between baseline and both the learning events for the group 1 animals and second learning event is found to be significant when compared with AIC and F-test (Table 2).The decrease in β, inverse of the clustering distance is indicative of increased clustering of spines. From our fits we estimate that learning extends the cooperative/clustering distance by 30% in group1 and 40 % in group 2 both compared with baseline. (F) The schematic of dendrites when scaled with their spine densities. Three dendrites with spine-density i) close to mean ii) higher than mean and iii) lesser than mean is shown. Higher density results in lower interspine distance thus contributing to false detection of cluster when averaged among dendrites with varying spine densities while lower density results in failed detection of cluster. Scaling measures spine location using mean interspine distance as metric thus accounting for the spatial effects that can arise from variation in spine density. Thus, enabling sensitive detection of clustering. (G) and (H) are the ACF of scaled dendrites from group 1 and group2 animals (solid circles) across four sessions and their fit (solid lines) to Eq. 4 stated in text. Comparison of the fit parameters(I-N) show thatRβ, the inverse of NCD decreases significantly(~30%) (Table-3) following learning. These, changes are seen only when the animals learn and administering CPP prevents these changes. The ACF from BL1 session of the group2 did not fit (Adj. R. Sq < 0.7) hence the parameters are not shown or considered. Also, we see that independent but correlating fraction (Rμ) proportional to spine density shows an increase only after novel learning. Levels of significance is indicated through asterisks (*) with each star representing an order of magnitude lower chance factor. n.s indicates a non-significant difference.

Previously, it has been shown that molecules such as H-Ras get activated at spines following stimulation and diffuse out along the dendrite before getting deactivated. Diffusion of such activated molecules has been shown to stabilise neighbouring spines *in vitro* and are hypothesised to facilitate temporal integration through memory allocation(15,36). We describe the spread of the activated molecules and their subsequent inactivation along the dendrites as diffusion-coupled reaction. Thereby, describing the concentration profile of activated molecules and hence the occurrence probability of spine along the dendrite. This when combined with Eq. 1 yields,

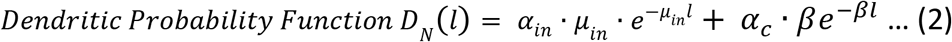

**Table 1A:** Post-hoc Analysis of ACF’s using Dunn’s Test. ACF till a delay of 125 pixels are compared across the four imaging sessions. We used non-parametric test, Friedman’s ANOVA to establish if the data is drawn from different distributions. (Chi-Sq: 73, p < 10-15).

|  | Sum Rank Diff | Z | Prob | Sig |
| --- | --- | --- | --- | --- |
| "BL1" "BL2" | -10 | 1 | 1 | 0 |
| "BL1" "L1" | -69 | 6.9 | 3.12015E-11 | 1 |
| "BL1" "L2" | -61 | 6.1 | 6.36411E-9 | 1 |
| "BL2" "L1" | -59 | 5.9 | 2.18101E-8 | 1 |
| "BL2" "L2" | -51 | 5.1 | 2.03792E-6 | 1 |
| "L1" "L2" | 8 | 0.8 | 1 | 0 |

**Table 1B:** Post-hoc Analysis of raw dendritic ACF’s and their comparison with ACF of the shuffled sequence (rand) using Dunn’s Test. Data set name BL1, BL2, L1 and L2 correspond to ACF from baseline1, baseline2, learning1 and learning2 sessions.

**Dunn's Test**
|  | Sum Rank Diff | Z | Prob | Sig |
| --- | --- | --- | --- | --- |
| "BL1" "BL2" | 37 | 0.81919 | 1 | 0 |
| "BL1" "L1" | -420 | 9.29896 | 3.97127E-19 | 1 |
| "BL1" "L2" | -352 | 7.79341 | 1.82627E-13 | 1 |
| "BL1" "BL1_Rand" | -218 | 4.8266 | 3.88872E-5 | 1 |
| "BL1" "BL2_Rand" | -228 | 5.048 | 1.25005E-5 | 1 |
| "BL1" "L1_rand" | -741 | 16.40602 | 4.84975E-59 | 1 |
| "BL1" "L2_rand" | -590 | 13.06282 | 1.50364E-37 | 1 |
| "BL2" "L1" | -457 | 10.11815 | 1.28523E-22 | 1 |
| "BL2" "L2" | -389 | 8.6126 | 1.99973E-16 | 1 |
| "BL2" "BL1_Rand" | -255 | 5.64579 | 4.60375E-7 | 1 |
| "BL2" "BL2_Rand" | -265 | 5.8672 | 1.24101E-7 | 1 |
| "BL2" "L1_rand" | -778 | 17.22521 | 4.81053E-65 | 1 |
| "BL2" "L2_rand" | -627 | 13.88201 | 2.27988E-42 | 1 |
| "L1" "L2" | 68 | 1.50555 | 1 | 0 |
| "L1" "BL1_Rand" | 202 | 4.47236 | 2.16616E-4 | 1 |
| "L1" "BL2_Rand" | 192 | 4.25095 | 5.9602E-4 | 1 |
| "L1" "L1_rand" | -321 | 7.10706 | 3.31916E-11 | 1 |
| "L1" "L2_rand" | -170 | 3.76386 | 0.00468 | 1 |
| "L2" "BL1_Rand" | 134 | 2.96681 | 0.08425 | 0 |
| "L2" "BL2_Rand" | 124 | 2.74541 | 0.16922 | 0 |
| "L2" "L1_rand" | -389 | 8.6126 | 1.99973E-16 | 1 |
| "L2" "L2_rand" | -238 | 5.26941 | 3.83219E-6 | 1 |
| "BL1_Rand" "BL2_Rand" | -10 | 0.2214 | 1 | 0 |
| "BL1_Rand" "L1_rand" | -523 | 11.57941 | 1.46727E-29 | 1 |
| "BL1_Rand" "L2_rand" | -372 | 8.23622 | 4.9767E-15 | 1 |
| "BL2_Rand" "L1_rand" | -513 | 11.35801 | 1.8947E-28 | 1 |
| "BL2_Rand" "L2_rand" | -362 | 8.01481 | 3.08849E-14 | 1 |
| "L1_rand" "L2_rand" | 151 | 3.3432 | 0.02319 | 1 |

**Table 2:** Comparison of β parameter of Group 1 ACF across sessions using AIC and F-Test.

| Dataset Compared | Akaike's Information Criteria |  |  |  | F- Test |  |
| --- | --- | --- | --- | --- | --- | --- |
|  | AIC Index |  | Weight of being different | Reduction in Information Loss | F | Prob (p- value) |
| Group 1 $\beta$ | Same | Different | | | | |
| BL1BL2 | -4.26E+3 | -4.26E+3 | 2.84E-1 | 3.97E-1 | 6.73E-1 | 4.12E-1 |
| L1BL2 | -4.17E+3 | -4.18E+3 | 9.94E-1 | 1.72E+2 | 1.28E+1 | 4.03E-4 |
| L1BL1 | -4.24E+3 | -4.24E+3 | 9.32E-1 | 1.36E+1 | 7.66E+0 | 5.95E-3 |
| L2BL1 | -4.23E+3 | -4.23E+3 | 9.47E-1 | 1.79E+1 | 8.21E+0 | 4.42E-3 |
| L2BL2 | -4.16E+3 | -4.17E+3 | 9.95E-1 | 2.20E+2 | 1.33E+1 | 3.12E-4 |
| L2L1 | -4.17E+3 | -4.16E+3 | 2.23E-1 | 2.87E-1 | 3.95E-2 | 8.43E-1 |

| Dataset Compared | AIC |  |  |  | F - test |  |
| --- | --- | --- | --- | --- | --- | --- |
| Group 2 $\beta$ | Same | Different | Weight of being different | Reduction in Information Loss | F | Prob (p – value) |
| bl1bl2 | -4.81E+3 | -4.81E+3 | 2.69E-1 | 3.68E-1 | 2.62E-3 | 9.59E-1 |
| l1b1 | -4.71E+3 | -4.71E+3 | 0.272 | 0.374 | 0.0303 | 0.862 |
| l1b2 | -4.79E+3 | -4.79E+3 | 0.272 | 0.373 | 0.0266 | 0.870 |
| l2b1 | -4.79E+3 | -4.80E+3 | 0.987 | 76.4 | 10.6 | 0.00123 |
| l1l2 | -4.77E+3 | -4.79E+3 | 0.999 | 1.26E+3 | 16.3 | 6.57E-5 |
| l2bl2 | -4.89E+3 | -4.90E+3 | 1.000 | 6.28E+3 | 19.6 | 1.25E-5 |

**Table 3:** Comparison of scaled ACFs of group 1 and group2:

Group 1:
| Dataset Compared | AIC |  |  |  | F - test |  |
| --- | --- | --- | --- | --- | --- | --- |
| Group1 $\beta_R$ | Same | Different | Weight of being different | Reduction in Information Loss | F | Prob |
| BL1BL2 | -5.42E+2 | -5.40E+2 | 2.67E-1 | 3.64E-1 | 6.14E-1 | 4.38E-1 |
| L1BL1 | -516 | -518 | 0.735 | 2.77 | 4.39 | 0.0420 |
| L1BL2 | -535 | -535 | 0.474 | 0.901 | 2.26 | 0.140 |
| L2BL1 | -515 | -526 | 0.997 | 365 | 14.8 | 3.80E-4 |
| L2BL2 | -537 | -547 | 0.995 | 186 | 13.3 | 7.11E-4 |
| L2L1 | -521 | -522 | 0.664 | 1.97 | 3.73 | 0.0598 |
| Group1 $\alpha_c$ | Same | Different | Weight of being different | Reduction in Information Loss | F | Prob |
| BL1BL2 | -5.43E+2 | -5.40E+2 | 2.15E-1 | 2.73E-1 | 1.05E-1 | 7.48E-1 |
| L1BL1 | -510 | -518 | 0.977 | 43.0 | 9.99 | 0.00284 |
| L1BL2 | -524 | -535 | 0.996 | 261 | 14.0 | 5.18E-4 |
| L2BL1 | -523 | -526 | 0.872 | 6.83 | 6.16 | 0.0169 |
| L2BL2 | -540 | -547 | 0.979 | 46.0 | 10.1 | 0.00266 |
| L2L1 | -525 | -522 | 0.214 | 0.273 | 0.102 | 0.751 |
| Group1 $\mu_R$ | Same | Different | Weight of being different | Reduction in Information Loss | F | Prob |
| | | | | $\exp((B-C)/2)$ | | |
| BL1BL2 | -5.41E+2 | -5.40E+2 | 3.32E-1 | 4.97E-1 | 1.17E+0 | 2.85E-1 |
| L1BL1 | -469 | -518 | 1.00 | 3.66E+10 | 78.9 | 2.26E-11 |
| L1BL2 | -481 | -535 | 1.00 | 4.02E+11 | 91.3 | 2.68E-12 |
| L2BL1 | -514 | -526 | 0.997 | 381 | 14.9 | 3.64E-4 |
| L2BL2 | -537 | -547 | 0.995 | 187 | 13.3 | 7.09E-4 |
| L2L1 | -499 | -522 | 1.000 | 1.20E+5 | 30.2 | 1.88E-6 |

Group 2:
| Dataset Compared | AIC |  |  |  | F - test |  |
| --- | --- | --- | --- | --- | --- | --- |
| Group2 $\beta_R$ | Same | Different | Weight of being different | Reduction in Information Loss | F | Prob |
| | | | | $\exp((B-C)/2)$ | | |
| BL1BL2 | -3.66E+2 | -3.64E+2 | 2.88E-1 | 4.04E-1 | 1.09E+0 | 3.04E-1 |
| L1BL1 | -378 | -377 | 0.435 | 0.770 | 2.23 | 0.146 |
| L1BL2 | -369 | -366 | 0.183 | 0.224 | 0.0940 | 0.761 |
| L2BL1 | -376 | -381 | 0.905 | 9.50 | 7.06 | 0.0125 |
| L2BL2 | -369 | -368 | 0.420 | 0.723 | 2.12 | 0.156 |
| L2L1 | -385 | -384 | 0.353 | 0.544 | 1.61 | 0.214 |
| Group2 $\alpha_c$ | Same | Different | Weight of being different | Reduction in Information Loss | F | Prob |
| | | | | $\exp((B-C)/2)$ | | |
| BL1BL2 | -3.67E+2 | -3.64E+2 | 1.97E-1 | 2.45E-1 | 2.42E-1 | 6.27E-1 |
| L1BL1 | -380 | -377 | 0.200 | 0.250 | 0.277 | 0.603 |
| L1BL2 | -369 | -366 | 0.187 | 0.230 | 0.139 | 0.712 |
| L2BL1 | -355 | -381 | 1.000 | 3.96E+5 | 36.9 | 1.13E-6 |
| L2BL2 | -363 | -368 | 0.941 | 15.9 | 8.13 | 0.00779 |
| L2L1 | -373 | -384 | 0.995 | 220 | 14.1 | 7.38E-4 |
| Group2 $\mu_R$ | Same | Different | Weight of being different | Reduction in Information Loss | F | Prob |
| | | | | $\exp((B-C)/2)$ | | |
| BL1BL2 | -3.67E+2 | -3.64E+2 | 1.77E-1 | 2.15E-1 | 2.79E-2 | 8.68E-1 |
| L1BL1 | -377 | -377 | 0.527 | 1.11 | 2.90 | 0.0991 |
| L1BL2 | -365 | -366 | 0.565 | 1.30 | 3.18 | 0.0848 |
| L2BL1 | -383 | -381 | 0.252 | 0.337 | 0.782 | 0.383 |
| L2BL2 | -361 | -368 | 0.971 | 33.1 | 9.72 | 0.00400 |
| L2L1 | -372 | -384 | 0.997 | 305 | 14.9 | 5.54E-4 |

**Table 4:** Comparison of new and lost spines ACFs of group 1 and group2: BetaR

| Dataset Compared | AIC |  |  |  | F - test |  |
| --- | --- | --- | --- | --- | --- | --- |
| Group3 $\beta_R$ | Same | Different | Weight of being different | Reduction in Information Loss<br>$\exp((B-C)/2)$ | F | Prob |
| BL1BL2 | -5.21E+2 | -5.19E+2 | 2.07E-1 | 2.61E-1 | 2.33E-2 | 8.79E-1 |
| L1BL1 | -494 | -495 | 0.655 | 1.90 | 3.67 | 0.0621 |
| L1BL2 | -494 | -496 | 0.704 | 2.38 | 4.10 | 0.0491 |
| L2BL1 | -467 | -513 | 1.00 | 1.09E+10 | 73.1 | 6.63E-11 |
| L2BL2 | -511 | -514 | 0.859 | 6.09 | 5.93 | 0.0190 |
| L1L2 | -495 | -492 | 0.204 | 0.256 | 0.00862 | 0.926 |

AlphaC
| Dataset Compared | AIC |  |  |  | F - test |  |
| --- | --- | --- | --- | --- | --- | --- |
| Group3 $\alpha_c$ | Same | Different | Weight of being different | Reduction in Information Loss<br>$\exp((B-C)/2)$ | F | Prob |
| BL1BL2 | -5.21E+2 | -5.19E+2 | 2.06E-1 | 2.59E-1 | 1.26E-2 | 9.11E-1 |
| L1BL1 | -496 | -495 | 0.396 | 0.657 | 1.69 | 0.200 |
| L1BL2 | -497 | -496 | 0.379 | 0.610 | 1.56 | 0.219 |
| L2BL1 | -516 | -514 | 0.369 | 0.585 | 1.47 | 0.232 |
| L2BL2 | -516 | -514 | 0.369 | 0.585 | 1.47 | 0.232 |
| L1L2 | -495 | -492 | 0.205 | 0.258 | 0.0184 | 0.893 |

MuR:
| Dataset Compared | AIC |  |  |  | F - test |  |
| --- | --- | --- | --- | --- | --- | --- |
| Group3 $\mu_R$ | Same | Different | Weight of being different | Reduction in Information Loss<br>$\exp((B-C)/2)$ | F | Prob |
| BL1BL2 | -5.10E+2 | -5.19E+2 | 9.88E-1 | 8.16E+1 | 1.14E+1 | 1.55E-3 |
| L1BL1 | -456 | -495 | 1.000 | 3.34E+8 | 58.3 | 1.57E-9 |
| L1BL2 | -447 | -496 | 1.00 | 4.85E+10 | 81.1 | 1.87E-11 |
| L2BL1 | -467 | -513 | 1.00 | 1.09E+10 | 73.1 | 6.63E-11 |
| L2BL2 | -492 | -514 | 1.000 | 7.24E+4 | 28.7 | 2.96E-6 |
| L1L2 | -479 | -492 | 0.998 | 654 | 16.2 | 2.24E-4 |

The above equation has two terms: first term describing the order that is brought about by the spine density and second term describing the modulation of stability due to diffusional exchange across the neighbouring spines. Eq.2 is arrived after normalising two terms to their respective fractions (refer to SI:Theory for detailed derivation). From Eq. 2 the spine distribution is characterised by clustered fraction (*α*_*C*_), the inactivation/influence length (*β*) and the spine density of independent spines (μ_in_). Given ACF’s ability to extract physical characteristics of quasi-periodic signals, we use ACF to extract the functional characteristics of clustering. The ACF for the above distribution is given by,

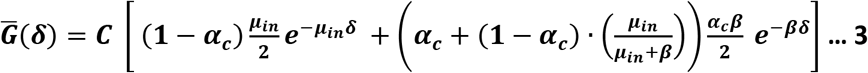

This enables us to characterise the changes in the spatial organization of all the spines and not just new spines in isolation. The autocorrelations thus obtained as a function of behavioral training (open circles in Fig. 2B for Grp 1 and Fig. 2C for Grp 2) start from a non-zero value at delay of one pixel and then progressively decay, converge to same value and ultimately decay to near-zero values. The correlation of the initial region (from delay 1 to 125) is found to be significantly different (Z > 5, p < 10^−6^) between learning and baseline conditions in the group that received saline (Grp 1) during first training (Friedman Test). On the other hand, when the memory for the first learning is blocked using CPP (Grp 2), we see that the correlation is higher only after second learning. Before interpreting these results in terms of changes in spatial organisation of spine in a dendrite, we investigate if this could be a reflection merely of altered spine densities. Given a mean spine density, organising these spines in one-dimensional space could give rise to non-zero autocorrelation. We argued if the correlations were to come from spine density and not due to spines being in specific locations, then shuffling their placement/order in the sequence should not alter the ACF. On the contrary, when we computed the ACF after random shuffling of the sequences in silico without altering spine density, we see the ACF (Fig. 2: dashed lines in 2B and 2C) to have a significantly (Z > 2.5, p <0.01) lower correlation in all the four cases. We suggest that the above characteristic of ACF (correlation greater than that of randomised dataset) could be taken as a signature of spine clustering. We note that this observation is completely independent of any proposed model.

Next, we quantitatively compare across different functional manipulations. First, we verify if such a model is consistent with the observed distribution of spines. We observe that Eq. 3 data fits (solid lines in Fig.2B and Fig.2C) very well to Eq. 3 (Adj. R-square > 0.9). Thus, we proceed to fit the ACF data and determine the functional characteristic β (across groups, Fig. 2D and 2E). We find that for both groups, β is not significantly different between the two baseline sessions (β BL1 = 1.033 ± 0.090 and BL2 = 1.121 ±0.102) using Akai’s Information Criterion (AIC) test. However, in Grp 1, whereas β is significantly low compared to baseline, first learning decreases β significantly (Table 2, F > 7, p <5E-3) while second learning does not cause any further decrease, even though β is still significantly low compared to the baseline (Fig. 2D). Thus, we interpret that learning is accompanied by significant increase in interspine cooperative distance or in other words cluster length.

Consistent with our interpretation in Grp 2 we see a significant decrease in β (Fig. 2E) only following second learning as learning is inhibited during the first session due to CPP. Despite such differences in β the clustering fraction(α_c_) does not change across groups (SFig. 2) although spine density has changed. Further it is useful to measure the clustering size as a fraction of average interspine distance. To achieve this, we scaled the dendrites with their spine density to measure the position of a spine in units of the average interspine distance. Such scaling decouples the spine density changes from clustering induced changes (Fig. 2F). In such a case, the ACF(Eq.3) is modified to

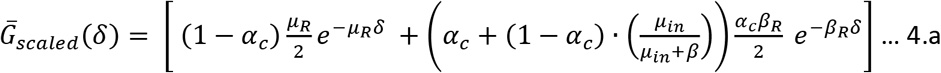

Which can be simplified, *for β* ≫ *μ*_*in*_,

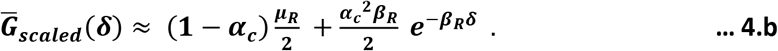

Using Eq. 4b above we can now measure the inactivation length in terms of interspine distance and clustered fraction. The ACF obtained after scaling, fits well (Adj R-sq > 0.94) to Eq. 4 (Fig. 2G, solid circles) yielding a measure of clustered fraction and clustering distance as a fraction of interspine distance (Normalised Clustering Distance (NCD)) (Fig. 2G-N). Consistent with our earlier observation, the NCD following first training event increased in Grp 1(bar graph, Fig. 2I) but not in Grp 2 and clustered fraction (α_c_) increased upon learning in both the groups (Figs. 2K and 2M). The ACF of the Grp 2 animals that underwent context A training in presence of CPP (shown in black, red, blue and olive in 2H,2J,2L and 2N) are not significantly different from the ACF obtained from pre-training images.

Taken together we interpret that i) learning, both in presence as well as in absence, of NMDAR activation results in altered organisation of spines and increases clustering ii) and such an increase is prevented when learning is blocked by administering CPP.

Next we probed, if the addition of new, or loss of old spines in response to new learning is clustered. We study this by constructing two dendritic sequences i) with only new spines and ii) with just lost spines. Operationally, we define spines present in that session but absent till then as new spines and lost spines are spines absent in the current session but present in the previous imaging session. Thus, ACFs are generated for each group by comparing the pairs, BL1-BL2, BL2-L1 and L1-L2 (Fig. 3). As expected, addition and removal of spines before training is random, since the baseline ACF for these cases are comparable to the shuffled ACF and fit Eq. 4b poorly (Adj R-Sq <0.7). In comparison, following novel learning events (L1 in Grp 1 and L2 in Grp 2) the new spine ACF shows a significant increase in the NCD (decrease in β) (Fig. 3E) as well as in the clustered fraction(α_c_)(Fig. 3F). For comparison, the clustering of the added spines is also shown using the conventional cumulative frequency distribution (CFD) of nearest neighbour distances for new spines (SFig. 5). However, such an increase in clustering in added spines is absent for related learning, suggesting clustered addition of spine is seen during new learning but not during related learning. Contrastingly, lost spines show (Fig. 3B, 3E-G) a clustering (increased fraction and NCD) only after related learning and not after new learning in Grp1, suggesting that the selective loss of spines seen could be representing the encoding of related memories. Intriguingly, animals from Grp 2 show clustered loss (increased NCD and α_c_) during first training event even though the animals were under CPP blockade and not during second training event where the CPP blockade was removed. This presents one of two possibilities: i) acquiring information related to prior learning is characterised by selective spine loss, though it may not be sufficient to elicit a behavioral response on its own or ii) NMDAR blockade results in clustered loss of spines.

**Figure 3:**
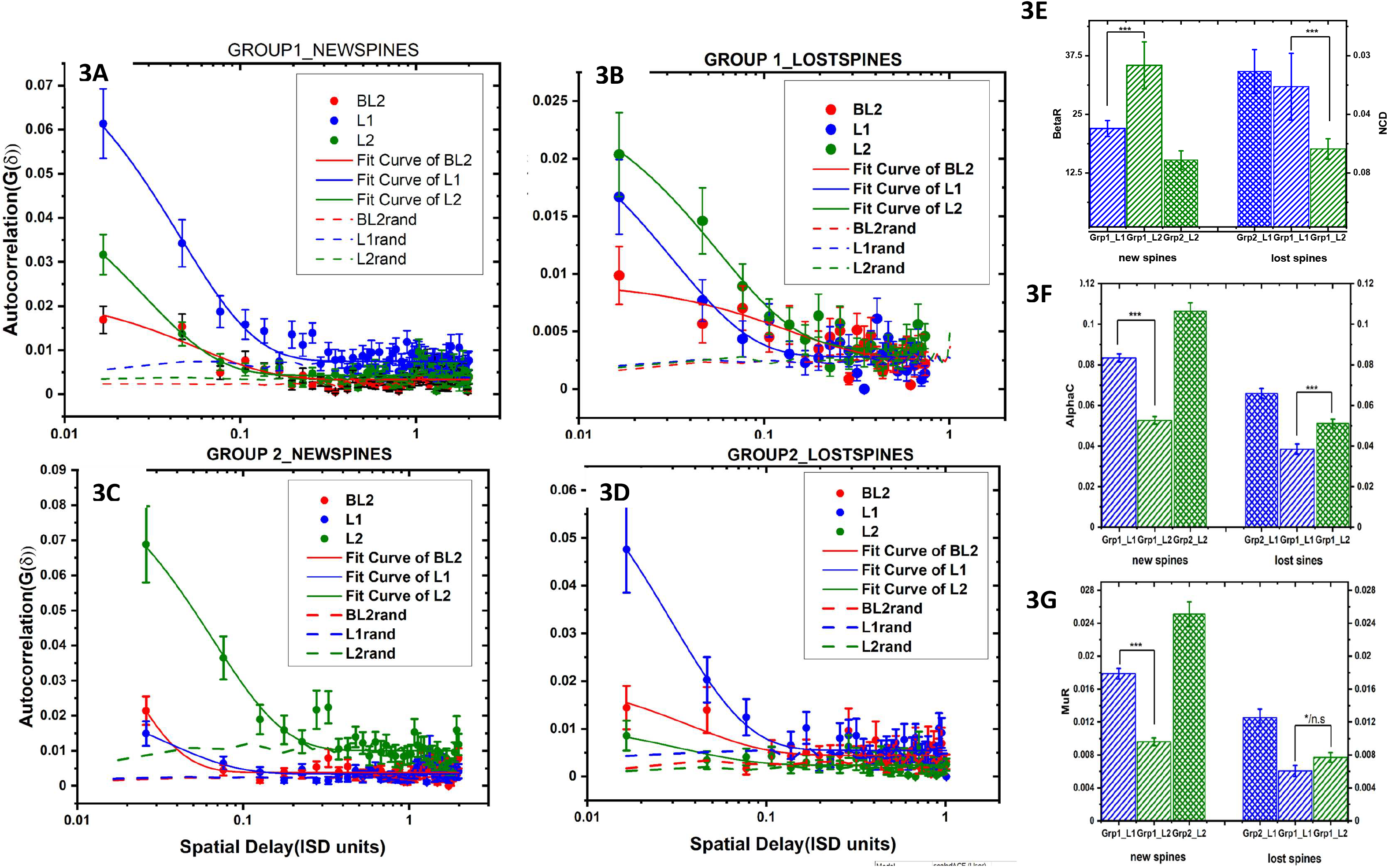
New spines show clustered addition during novel learning while clustered spine loss is dominant during related learning. (A) The ACF obtained from dendritic sequence of just new spines detected across imaging sessions of group 1 are shown as solid circles. The solid lines are the fits. (B)Plot of the ACF obtained from sequence that contained only the lost spines (solid circles) and their fit (solid line). Dashed lines in both these graphs are the ACF obtained after shuffling. (C) and (D) are the corresponding plots of ACF’s for the group2 mice. (E)-(G) Bar graphs comparing the fit parameters (β_R_, α_c_ and μ_R_) for new (left) and lost (right) spines. Fit parameters of first(blue) and second(olive) learning sessions of group 1(slanted lines) and group2(hatched) show addition of new spines exhibit declustering (increase in β_R_ and/or decrease in α_c_) and spine loss is clustered during related learning while regular learning results in clustered addition of new spines. We used adjusted R squared to assess the quality of fit. Adj. R. Sq. < 0.7 is considered a poor fit. We interpret the lack of good fit as lack of correlation and/or absence of clustering, thus new spines added or spine loss during the baseline sessions, spine loss following first learning all lack of correlation and hence we interpret as clustering is absent.

If clustering and loss of spines are characteristics of related learning, then animals undergoing regular training without NMDAR blockade in both sessions, should show similar clustering. Therefore, we ran a group of mice in a similar behavioral design (Saline infusion, Fig. 4A) except now the animals receive saline before both training sessions. Consistent with our hypothesis the ACF derived from all spines (Fig. 4B) and scaled dendrites (Fig. 4C) show increased NCD following new learning (Fig. 4E) without any further increase following related learning. New spines are added in a cluster as *α*_*c*_ increases only following new learning and not after related learning (Fig. 4H-J). Contrarily, spine loss in this group still shows an increase in NCD and *α*_*c*_ following related learning and not after new learning (Fig. 4K-M). This supports our hypothesis that it is indeed the acquisition of information related to prior learning that causes clustered loss of spines.

**Figure 4:**
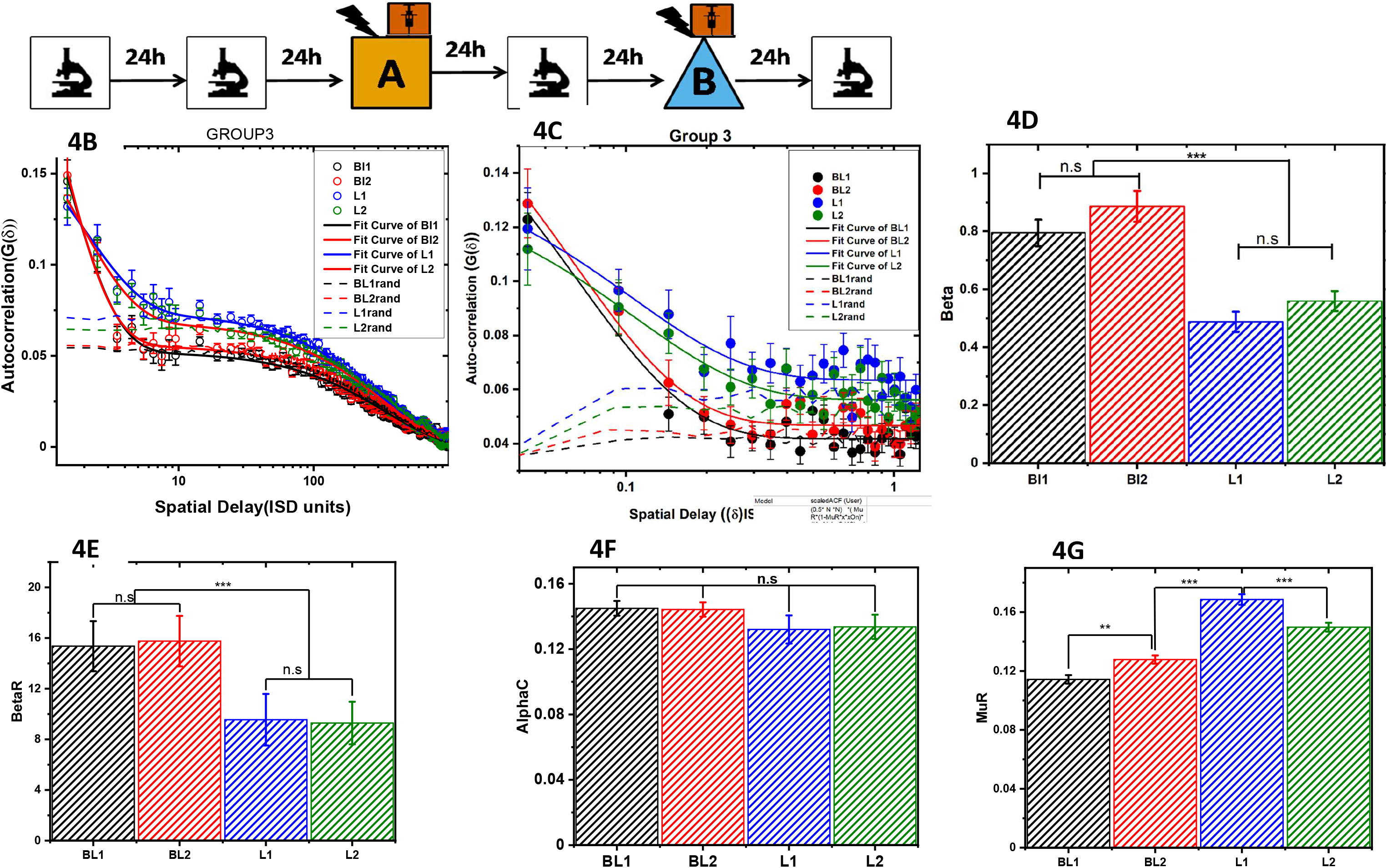
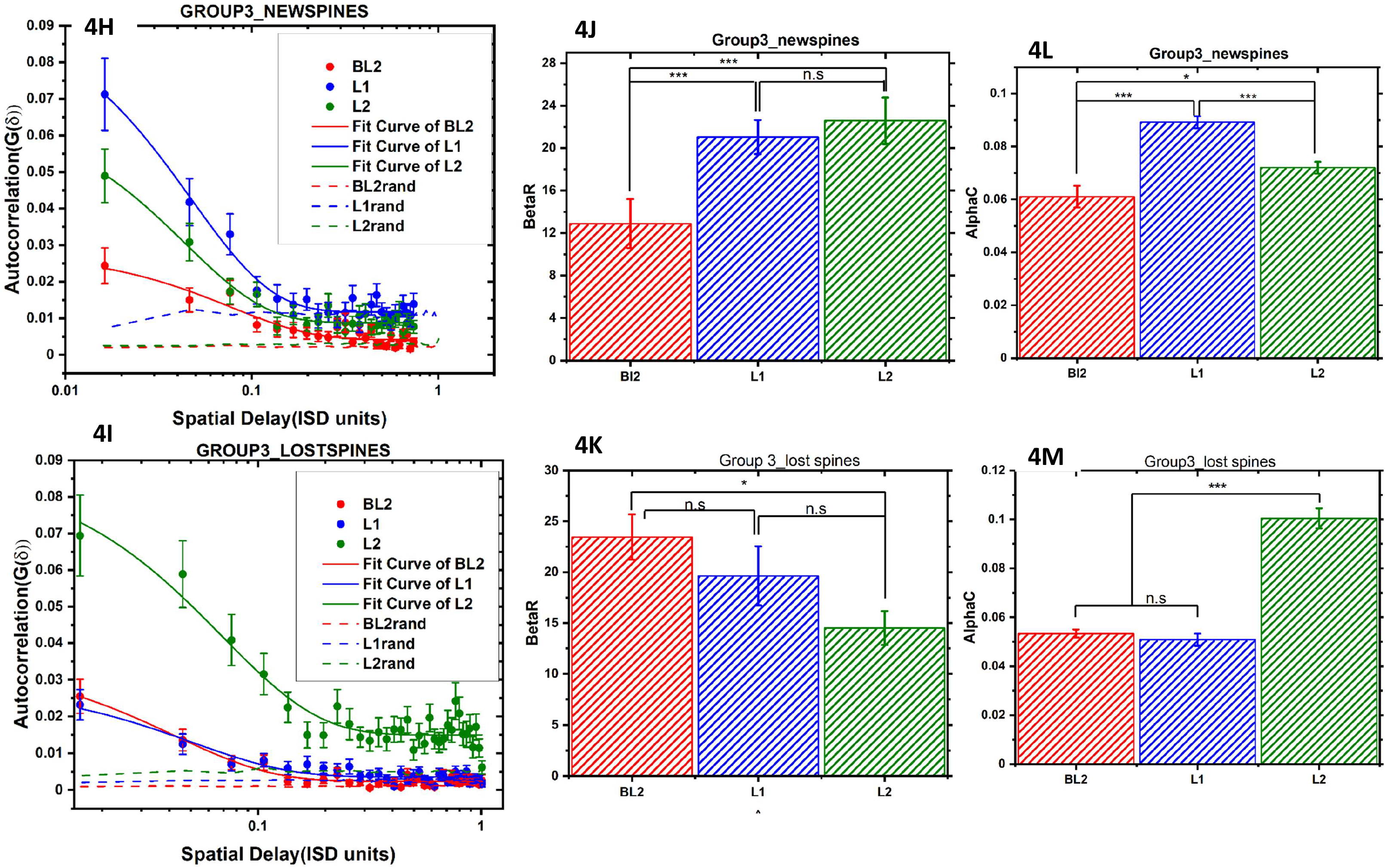
Related learning in absence of NMDAr blockers shows clustered loss of spines. (A) Imaging scheme used for animals in group3. The animals are trained in two contexts in presence of saline unlike the other two group of animals. (B) Solid circles are the ACF of raw dendritic sequence obtained from 70 dendrites across four imaging sessions viz baseline 1(BL1, black), baseline2(BL2), learning1(L1) and learning2(L2). Solid lines are the fits and dashed lines are the ACF of the shuffled dendritic sequence. (C) The scaled ACF (solid circles) and their fits (solid lines) obtained during four imaging sessions along with the ACF of the shuffled sequence are shown. (D)-(G) Comparison of the fit parameters obtained from (B) and (C) are shown as the bar graphs. (H) and (I) shows the ACF of the new and lost spines respectively in this group. The fit parameters and their comparison are shown in (J)-(M) with red, blue and green bars corresponding to baseline, novel learning and related learning sessions. Table-5 lists the AIC and F Test statistics of these parameters. The goodness of fit, estimation of statistical significance are done as explained earlier using Adj. R.Sq and AIC. Table 5 in SI provides a detailed listing of these statistic.

As related learning is independent of NMDAR activation and prior learning is a prerequisite for such independence, we reasoned that the structural correlates of the two learning events could be spatially linked. If clustered spine loss represents related information, then such loss should presumably occur in proximity to the spatial location of the spines added during formation of original memory. Thus, cross correlating the dendritic sequence of lost spines following related memory formation with that of new-spine sequence corresponding to original memory should uncover the spatial relationship that might exist between these phenomena. Indeed, both group 1 and group 3 cross-correlation functions (CCF) (Fig. 5A and 5C) show a characteristic clustering with longer NCD and higher clustering fraction. As expected of Grp 2 animals, their CCF do not exhibit any such clustering. Together we conclude that clustered loss of spines observed for the second learning is spatially linked to the clustered gain due to the first novel learning.

**Figure 5:**
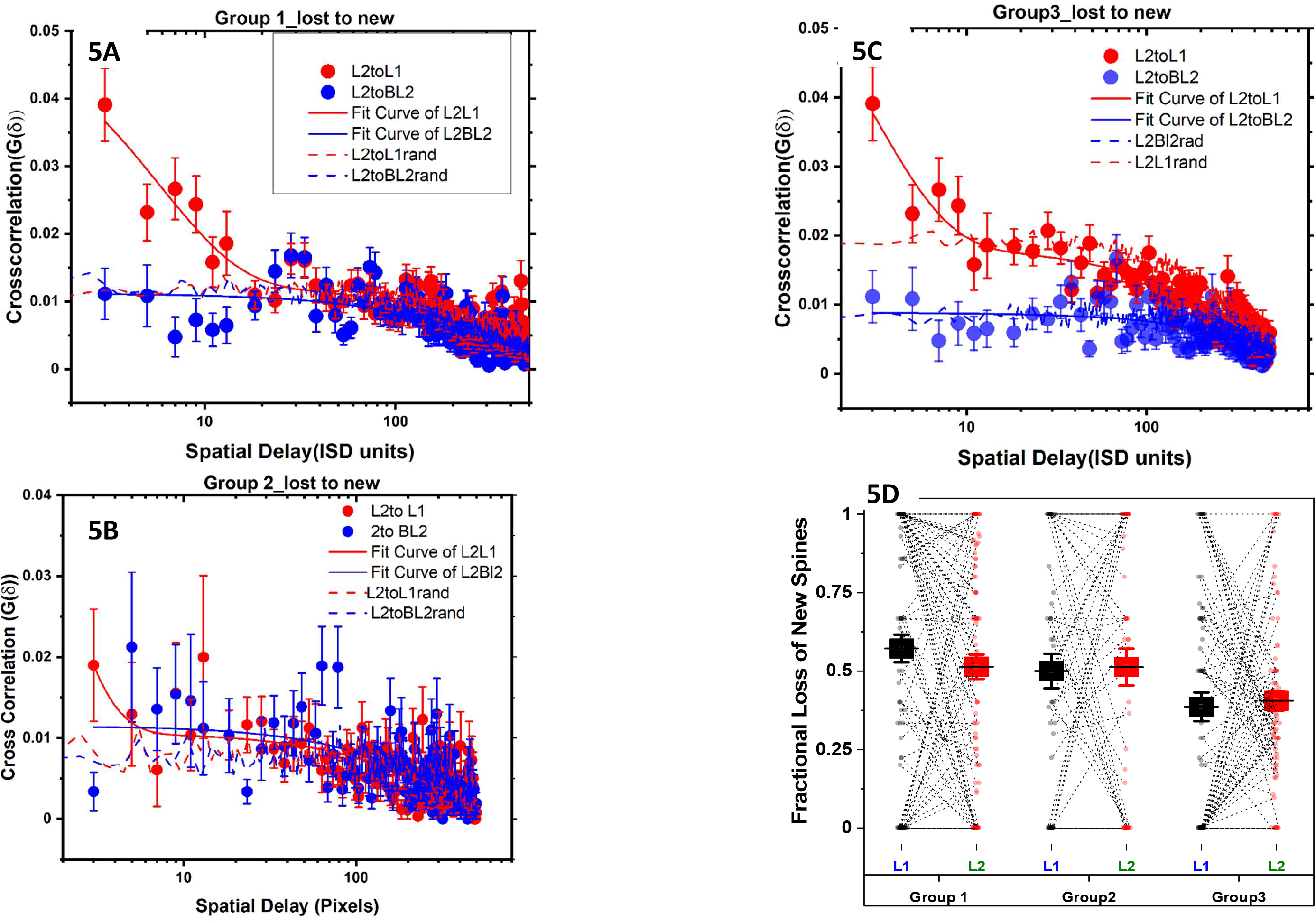
Cross-correlation of new spines formed during formation of initial memory with spines lost during related learning establish it is selective and clustered loss rather than gain that represent related memory. (A) Cross-correlation of spines lost during learning of related context and new spines formed during acquisition of initial context (red solid circles) show a prolonged decay compared to cross-correlation of new spines from baseline (black solid circles). The later CCF is similar to dashed lines that represent the cross correlation obtained after shuffling the order of spine locations. The solid lines are the fit for group1, (B) group2 and (C) group3. Comparing the CCF from all the three groups and across different sessions reveal that loss of newly formed spines is spatially correlated only in the case of related learning irrespective of NDAr blockade.(D) Mean fractional loss of the new spines formed after baseline (BL2) and lost after first-learning (L1) (black square) is compared with those formed after first-learning(L1) and lost after second learning(L2) (red square) for all three groups ((n > 100 for group1, n > 70 for group2 and n > 90 for group 3). The fractional loss were not different between sessions (ANOVA, F = 0.2, p >0.65). Existence of spatial correlation despite similar fractional loss of new spines suggests it is the relationship among the memories rather than the spine half-life that gives rise to spatially correlated loss.

**Table 5:** Comparison of ACFs of group 3:

| Dataset Compared | AIC |  |  |  | F - test |  |
| --- | --- | --- | --- | --- | --- | --- |
| Group3 $\beta$ | Same | Different | Weight of being different | Reduction in Information Loss<br>$\exp((B-C)/2)$ | F | Prob |
| BL1BL2 | -5.63E+3 | -5.63E+3 | 5.06E-1 | 1.02E+0 | 2.02E+0 | 1.56E-1 |
| L1BL1 | -5.66E+3 | -5.68E+3 | 1.000 | 1.14E+5 | 25.5 | 6.26E-7 |
| L1BL2 | -5.65E+3 | -5.69E+3 | 1.000 | 3.45E+7 | 37.5 | 1.92E-9 |
| L2BL1 | -5.69E+3 | -5.71E+3 | 0.999 | 736 | 15.2 | 1.12E-4 |
| L2BL2 | -5.69E+3 | -5.71E+3 | 1.000 | 1.20E+5 | 25.6 | 5.92E-7 |
| L1L2 | -5.77E+3 | -5.77E+3 | 0.483 | 0.934 | 1.83 | 0.176 |

Increase in correlation could arise from the transient nature of new spines since cross correlation can arise due to new spines that have short lifetime. To rule out this possibility, we compare the fractional loss of the new spines across two consecutive sessions (BL2/L1 and L2/L1) in all three groups (Fig. 5D). Two-way ANOVA with Groups and Sessions as factors showed that fractional loss is not different across sessions in all three groups (F = 0.2, p > 0.65 (for session)). Together with the fact that we see cross correlation differences only following related learning (Group 1 and Group2), and absent in group2 where the fraction loss of new spine is not different from Group 1, suggests that the correlated loss is specific to related memory formation. Additionally, we compare CCF of lost spines during the related memory formation with that of the new spines formed during baseline BL2 (Fig. 5A and 5C, solid blue line). These CCFs are comparable to correlation obtained from random shuffle (Dashed blue line) indicating that correlated loss is indeed due to refinement of the clusters specific to related memory. Further, we also test for similar correlations during the new memory formation by comparing the CCF following related learning with that of new memory (SFig. 3). While new memory exhibits CCFs with higher correlation than random shuffle, the extent of such clustering is significantly lesser than related memory formation as characterised by fits. These results suggest that clustered and correlated loss of dendritic spines characterize encoding of related memory. Thus using our new method we are able to show that spine loss does not necessarily mean a memory loss and infact it is common when encoding related memory.

In this process we formulate a method for probabilistic representation of spines in a dendrite and its relation to diffusional exchange between spines (Theory section of SI). Our formulation agrees well with the experimentally observed data obtained through high resolution longitudinal in *vivo* imaging of mice. This enabled us to show that dendrites do possess a stable clustered organisation of spines even before any behavioral training possibly, resulting from previous lifetime experiences. We estimate that about 15% of the spines show clustered organisation with a clustering distance of ~1/10^th^ the interspine distance (1/β_R_ of group 1 baseline) even at the basal state. Till now it has not been possible to detect such clustering as there is no known method for measuring absolute changes. Through ACFs, we measure that learning further extends the clustering distance 3 times from the baseline. We see, while new memories are characterised by clustered addition apart from other correlates of memory, related memory acquisition results in selective and spatially clustered pruning of the newly acquired spines. Thus, substantiating our claim that spine loss is an underlying feature of memory encoding.

## Acknowledgements

Grants: This work is funded by grants to BJ from SERB (EMR/2017/004155), DBT IISc Partnership, Ramanujan Fellowship, and Pratiksha Trust. MP is a recipient of CSIR fellowship 09/079(2697)/2016-EMR-I.

We would like to thank Prof. K P Yogendran, IISER Mohali for suggesting wait time distributions and for reading through the drafts and providing vital corrections, Prof. S Ray, CNS IISc for suggesting ACF of shuffled dendrites and Prof. S Maiti for discussions on ACFs and strong encouragement.

## Author Contributions

SKS, MP and BJ designed the experiment, SKS and MP performed the experiments, SKS and YK performed the Image and Data Analysis, BJ and SKS developed the theory analysed the data and BJ, SKS and MP wrote the paper.

## Theory: Organisation of the Spines in Dendrites

### Generating dendritic representation and its ACF

Consider a dendrite segment in RSc of length (“l”) with average spine density of (ρ = # of spines/length). Given an image of this dendrite, a sequence of 1’s and 0’s indicating the presence and absence of spines for a given length (measured in number of pixels) can be generated to represent the dendrite. A spine on a dendrite is treated as a reference point and the measurement as an origin about which the length of occurrence of other spines are measured. Thus, for a dendrite “i” we can write,

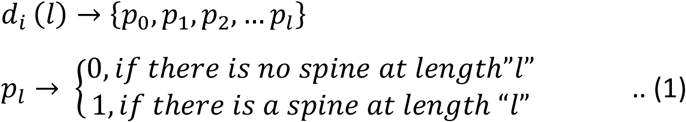

with *N*_*i*_ being the total number of spines present in the dendrite “i”.

In order to detect if there are any ordering among the spines, we estimate the 1D spatial autocorrelation

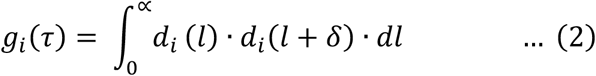

Given that there are no drastic changes in mean spine density across the dendrite segments in the RSc and distribution of the spines in each of the dendrites are independent we can exchange the order of integration and averaging. Essentially we require the cross correlation term di(l)dj(k) = 0. The average autocorrelation *G*(*τ*) is then obtained by averaging over the dendrites,

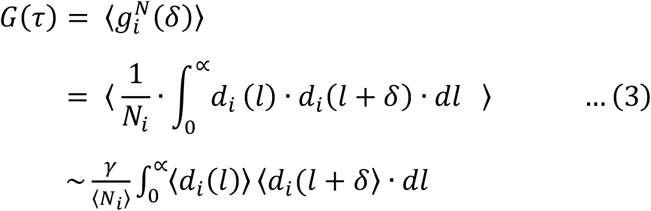

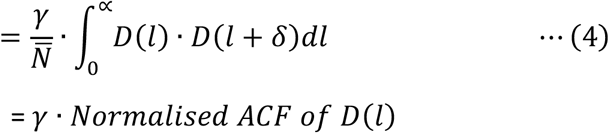

The “D(l)” being the normalised average representation of all the dendrites. This represents the probability density function corresponding to the spine distribution in a dendrite. γ is the constant that characterises the variation in the distribution of number of spines across different dendrites.

### Modelling of D(l)

We recognise there could be correlation corresponding to different spatial scales and given our optical resolution we restrict all our analysis to spatial delay of one pixel or more. Among this fraction of the spines whose spatial correlation persists beyond a pixel we realise that the mere presence of “N” number of spines in a stochastic manner on a length of “l” in an independent manner would impose a pattern and we call it as independent fraction. Being distributed stochastically with a mean density of “*μ*_*in*_” we can write the occurrence probability of such an “event” as a Poisson process. For a Poisson process the number of occurrences in a given time interval is independent of each other and are described by a Poisson distribution. However, the time interval between occurrences of events, termed as wait time (time one needs to wait before a desired event occurs), is described by the wait time distribution. For a first order Poisson they correspond to Erlang distribution. The wait time in this scenario corresponds to inter spine distance (distance we have to travel along the dendrite before we get another spine). Given that we measure the distance in a dendrite with respect to a spine(SFig. 1T), the probability of finding a spine at a distance “l” can be equated to probability of finding the inter spine distance “l”. Thus from the above arguments, the contribution of independent fraction to the probability density function of dendrite is given by,

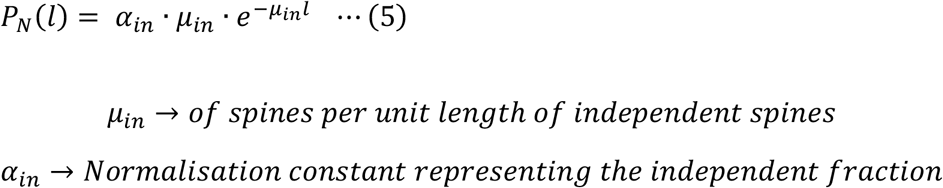

**SFig. 1T:**
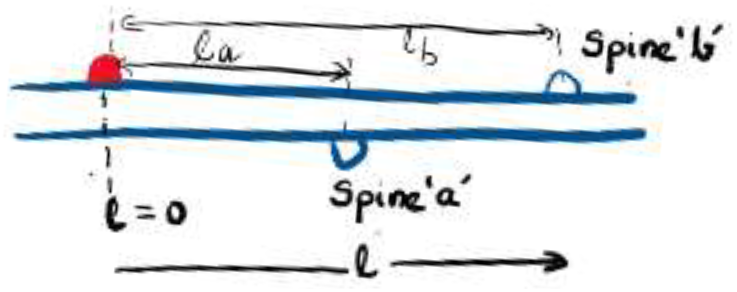
Co-ordinate system and lengths used to measure inter spine distances. Red spine is the point of origin, *l*_*a*_ and *l*_*b*_ represent the distances of spine “a” and “b” from origin.

There is another independent factor that affects the probability. Previously Harvey et al., have shown that diffusion of small molecules such as H-Ras from an activated spine biases the stability of neighbouring spines. Further, it has also been found (A. Govindarajan et al) that neighbouring spine can alter the stability through the diffusion of excitability related proteins (ERPs) and plasticity related proteins (PRPs). In general, if the information from the neighbouring spine is reaching the neighbours through diffusion and if the information is attenuated along the dendritic length(ref. #) the concentration of such molecules along the dendrite at steady state 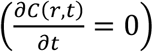 can be written as,

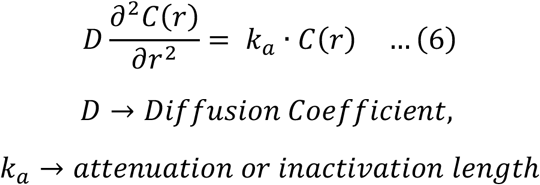

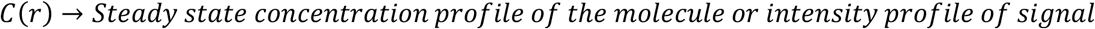

The solution for the above equation (6) can be found using the boundary conditions that the concentration would vanish at infinity and the concentration of the signal at l = 0 at steady state is given by 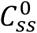.

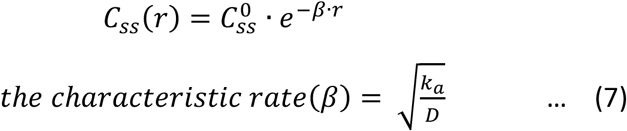

Note β is the inverse of the characteristic length over which the information/ material exchange and hence the correlation exists. We define a parameter, clustering distance NCD = 1 / β.

Eq. 7 describes the contribution of a spine in altering the stability of a neighbouring spine located at a distance “*r”*. In order to estimate the effect of such neighbouring spines at a dendritic position “l” we need to find the total contribution of all the spines that are present at different distances around “l”. Thus integrating such contribution across the dendritic length we can arrive at this contribution. However, we find that including the contribution from the entire length of the dendrite (no limit) is non-physical and leads to divergent solutions while restricting the contribution to a critical distance “*l*_*c*_” on either side of the spine at “l” provides a tractable solution. We will proceed to estimate the probability of finding a spine that is influenced by the activity of the neighbouring spines in this scenario(SFig. 2T). Red spine at *l = 0* defines the origin, red spine at “*l*” is where we estimate the influence of the neighbouring spines (green) present around red spine at “*l*”. We have shown the green spine at the the left to be at “x”and “*l*_*c*_” marks the critical distance. In such case the occurrence probability is given by, Occurrence probability of clustered spine at “l”

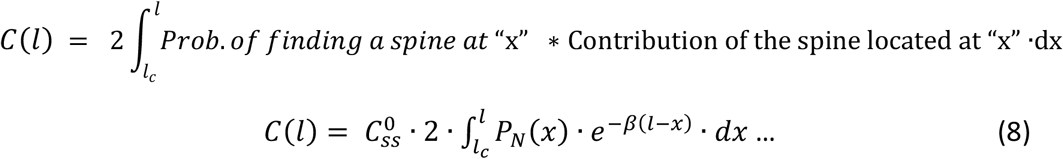

**SFig. 2T:**
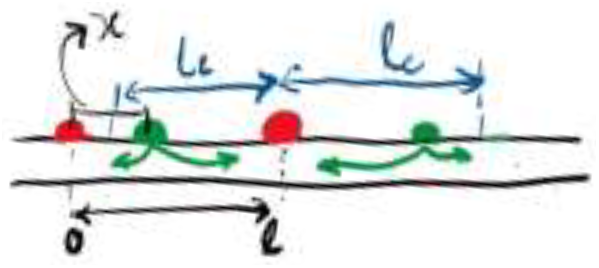
Estimating the influence from all the neighbouring spines (green).The diffusional contribution of spine at “x” is integrated over a critical length “*l*_*c*_”.

From Eq. 5 we can write,

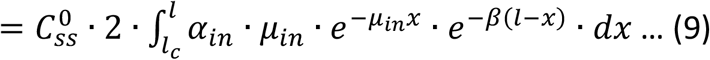

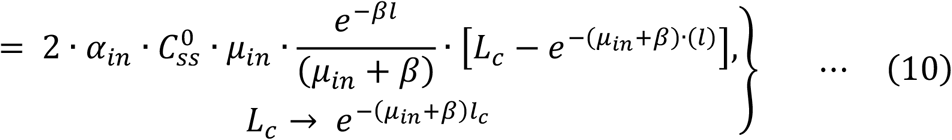

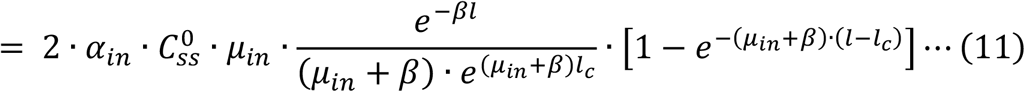

The constant 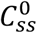 the number density of activated molecules generated in response to activity at the spine. This can be thought of a normalisation constant and determines the fraction of the spines that will be clustered/interacting.

At this point we consider two distinct models each differing in functions of the diffusing molecules, i) diffusing molecules merely stabilises ***existing*** synapses already present at “l” and ii) diffusing molecules not only can stabilise spines if present but can also ***create new*** if spine is absent at “l”. We found that the model described by (i) does not explain the observed data (substantially poor fit). Therefore we assume that the diffusing molecule also can create the spines.

In such a scenario,

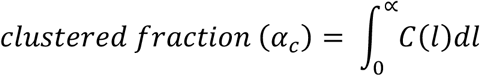

This implies

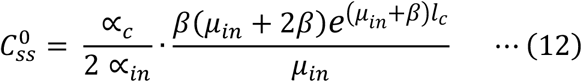

Using the above in Eq. 11 we have,

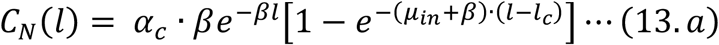

For the case 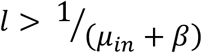 the above equation simplifies to

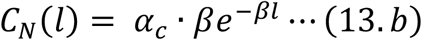

where *C*_*N*_(*l*) is the contribution of interspine interaction to normalised occurrence probability.

Now we can write the dendritic sequence D(l) which is the sum of the contribution from independent fraction PN(l) and clustering fraction CN(l). Thus, using Eq. 13.b, Eq. (5) and Eq. (4) we obtain the autocorrelation function as

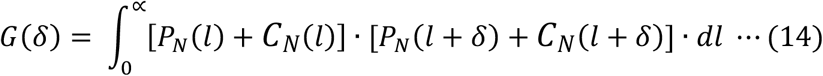

The autocorrelation has four different terms *I*_*PP*_, *I*_*CC*_, *I*_*PC*_ and *I*_*CP*_. Each of these terms can be evaluated individually

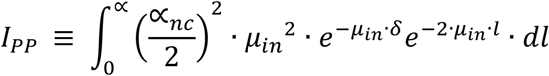

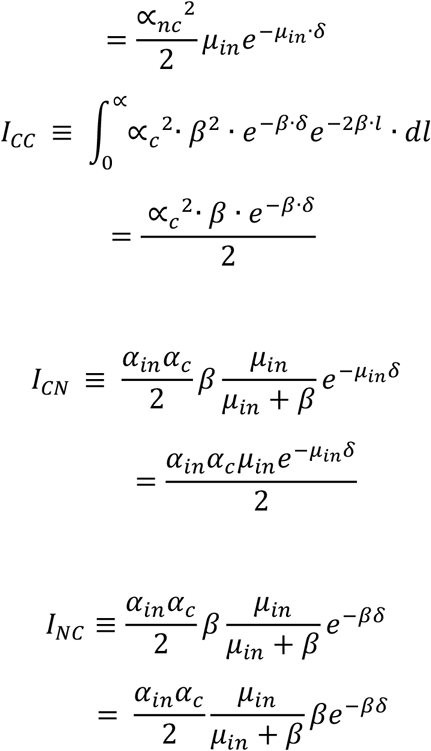

Using the above in Eq. (14) the autocorrelation for D(l) would be (14) the autocorrelation for D(l) would be

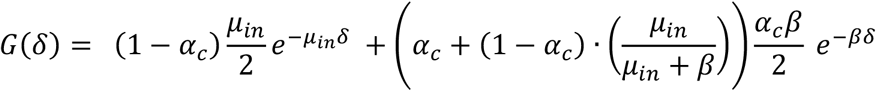

However, for average ACF as defined in Eq. (4), we have,

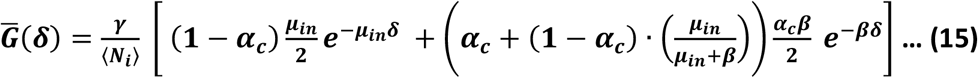

The above equation(Eq. 15) describes the ACF that can be used to describe and measure the extent of clustering from the physical parameters of inactivation length (1/β), spine density of independent spines(μ_in_), Fraction of interacting spines (α_c_).

**SFig. T3:**
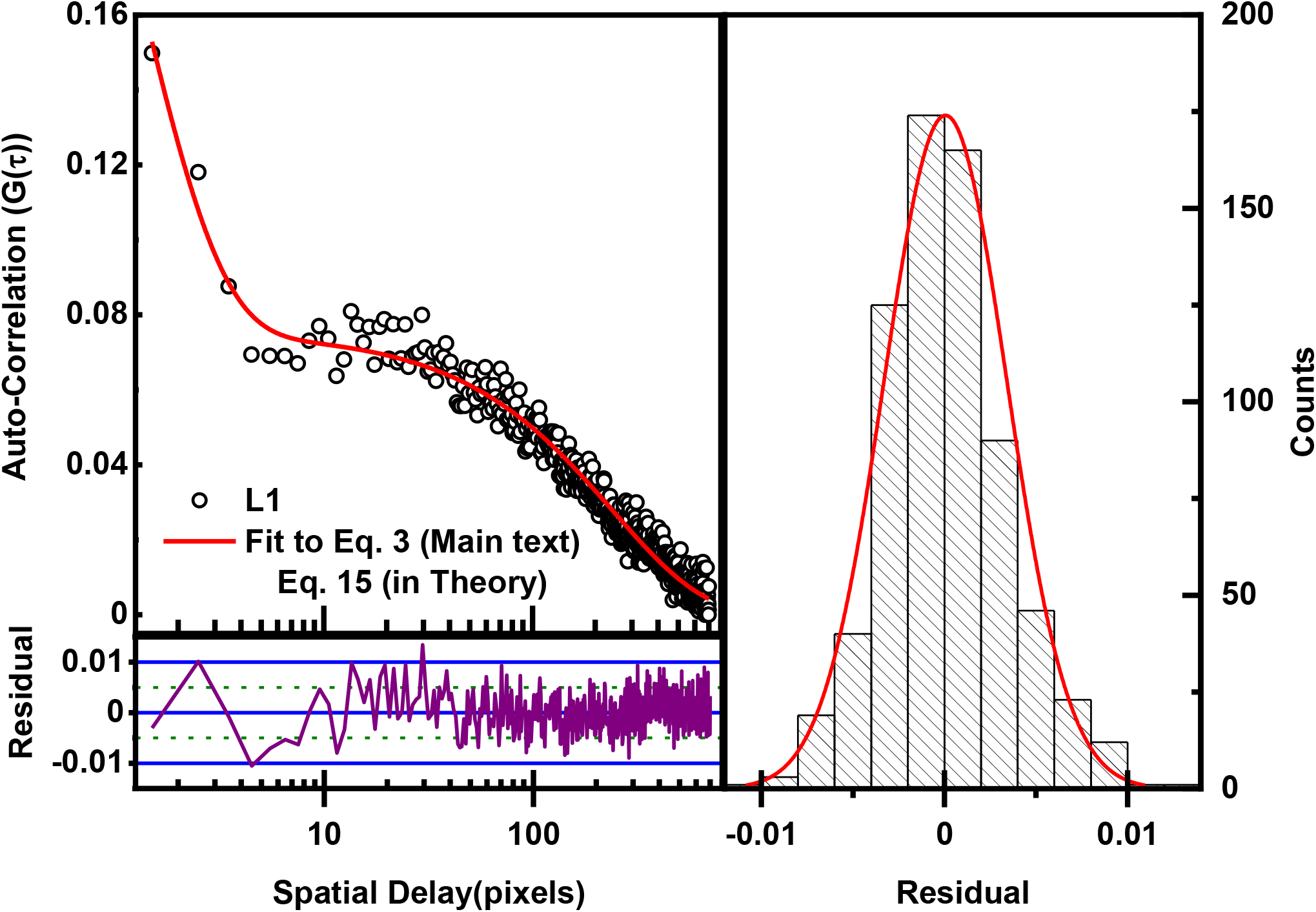
Black open circles on the top left panel are the scaled ACF data obtained from session L1 and the solid red line is the fit of Eq. 15 to this data. The residuals of the fit as a function of the spatial delay and the histogram of the residuals are shown in sperate panels. The even distribution of residual indicates a good fit (Adj. R Sq > 0.97). Error bars are not shown to better visualise the agreement of the data point and the fit curve.

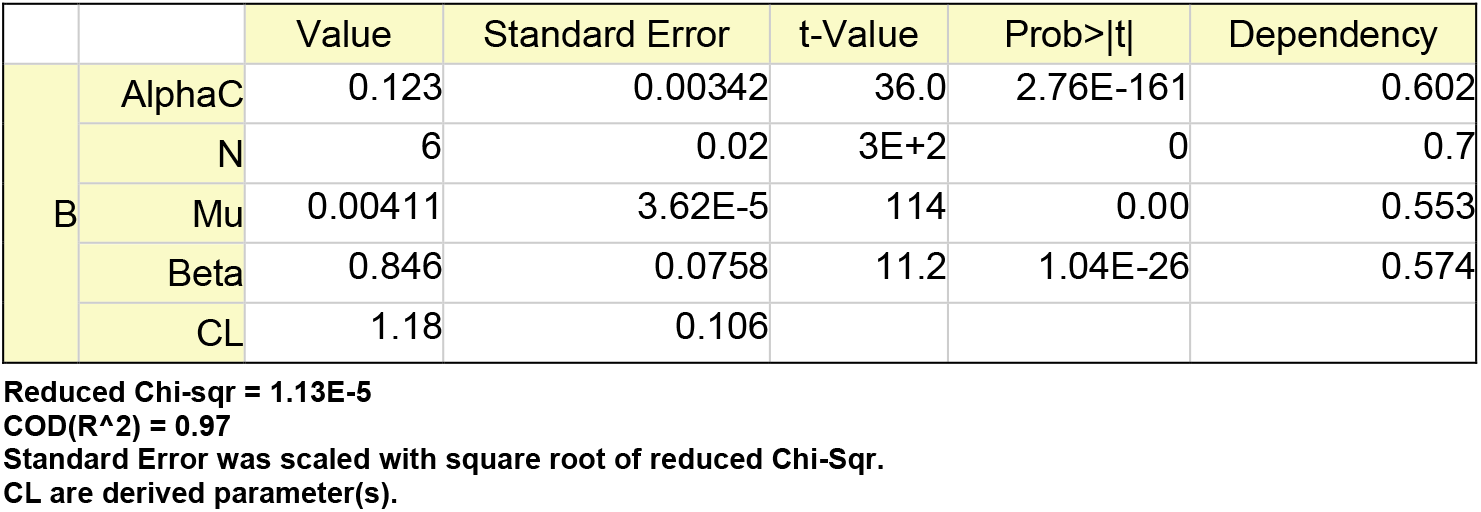

While the above equation provides a first analytical expression for characterising spine distribution. We note that this equation measures the inactivation length on laboratory frame agnostic to variation in spine density. Fraction of interacting spines, on the other hand strongly depends on spine density and variation in spine density can obscure the true measurement of α_c_. In order to address this we introduce a new scaled representation. In this representation the dendrites are scaled by the spine density such that all the dendrites have uniform spine density before ACF estimates. Thus we have 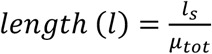, Thus, the scaled ACF can be written as,

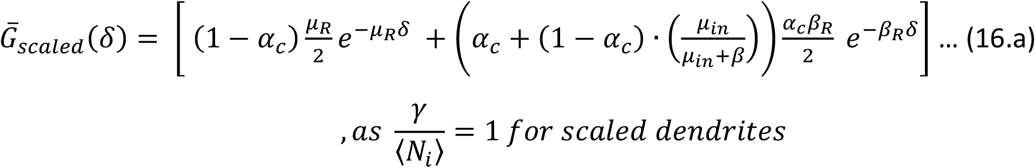

Further restricting the *δ* to around mean spine density, we can expand the first term and simplify as below:

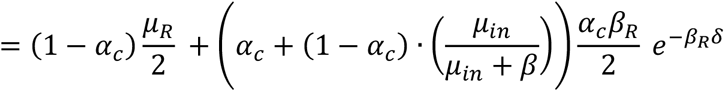

For β ≫ *μ*_*in*_ the above Eq. 14 further reduces to

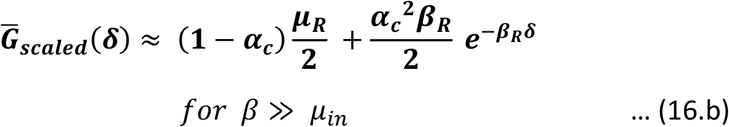

We use the above equation with α_c_, β_R_, and μ_R_ as parameters to fit and obtain the clustering fraction, inactivation length and mean independent spine density. We use the above equation with clustering fraction (α_c_), inactivation length (1/β_R_), and mean spine density of independent spine (μ_R_) as parameters from the fit.

**SFig. T4:**
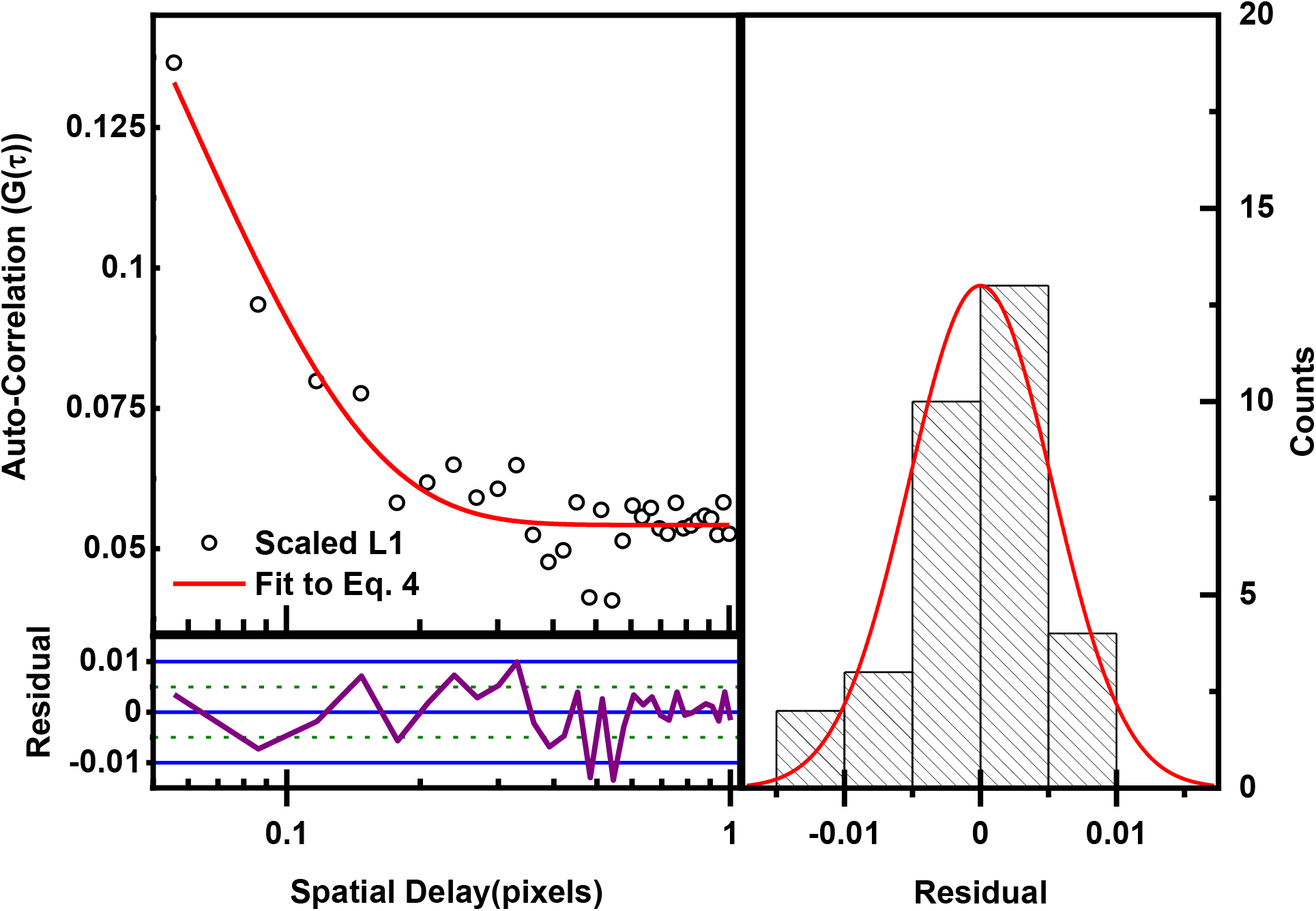
Black open circles on the top left panel are the ACF data obtained from session L1 and the solid red line is the fit of Eq. 16.b to this data. The residuals of the fit as a function of the spatial delay and the histogram of the residuals are shown in sperate panels. The even distribution of residual indicates a good fit (Adj. R Sq = 0.9). Error bars are not shown to better visualise the agreement of the data point and the fit curve.

Parameters
|  | Value | Standard Error | t-Value | Prob>\|t\| |
| --- | --- | --- | --- | --- |
| **AlphaC** | **0.155** | **0.00493** | **31.4** | **6.11E-24** |
| **BetaR** | **17.2** | **2.18** | **7.91** | **1.02E-8** |
| **MuR** | **0.128** | **0.00260** | **49.3** | **1.65E-29** |

Reduced Chi-sqr = 2.98E-5
COD(R<sup>2</sup>) = 0.91.
Standard Error was scaled with square root of reduced Chi-Sqr.

## Materials and methods

### Animals

Adult (3–8 months old) Male B6/129 F1 hybrid and Thy1-YFP-H (B6.Cg-Tg(Thy1-YFP)HJrs/J, Stock No: 003782) mice were used for the experiments as explained in the text. Mice were housed three per cage and given ad libitum food and water. Mice were maintained on a 12-h light/dark cycle and the behavioral tests were performed during the light phase of the cycle. All the mice were transferred to the holding room next to the contextual fear conditioning (CFC) room and handled for 5 mins for a week. The mice are habituated with the steps of the experiments such as transport process by individually taking the mice to the CFC room in the transport cage except for placing them in the context. The habituation was carried out for 2-3 days. All procedures are approved by the institute animal ethics committee.

### Contextual Fear Conditioning

Mice are trained and tested in custom made conditioning chambers (33 cm wide,30 cm high,30 cm deep) encased in sound-attenuated boxes (60 cm wide,50 cm high,80 cm deep). Inside chambers are made of clear acrylic. Every conditioning chamber consisted of CPU fan on the side wall, a stainless-steel grid floor (32 rods, each rod 4-mm diameter, 10-mm center to center spacing), a drop-pan and an overhead diffused LED-based light source. Video images were recorded using camera installed inside each box. The freezing response was measured using a custom plugin in imageJ called scoring assistant.

Context A features consisted of visible light, with fan noise and a single grid floor. The chambers and pans were cleaned with 70% ethanol prior to conditioning and recall test. In context B, visible light and fan was turned off and a translucent, triangular roof is placed inside the chamber. The grid floor contained steel rods in a staggered fashion, the chamber and pans are wiped with 50% isopropanol prior to conditioning and recall test.

During training, mice are placed in the context and allowed to explore for 1.5 min prior to shock onset. A mild foot shock of 0.75mA was delivered for 2 seconds. A minute after the shock, the mice are removed from the chamber and returned to their home cages.

### Injections

NMDAR antagonist CPP (Sigma-Aldrich, St. Louis, MO, USA) made by dissolving it in 0.9% saline. CPP was administered intraperitoneally 30 mins prior to training (10mg/kg, ~150μl).

### Surgery and Cranial Window Implantation

Our surgical procedures followed for cranial window implantation is adopted from methods previously described1. Mice are anesthetized using isoflurane and mounted on the stereotaxic instrument from David Kopf Instrument, USA (https://kopfinstruments.com/). Carprofen (5mg/kg) and dexamethasone(5mg/kg) are administered subcutaneously, and mice are kept warm using a monitored heating pad. After making an incision on top of the skull using intraaural distance as guide. The skin is retracted to the side exposing the skull. A region 4mm in radius is marked using stereotaxic coordinates (RSC: center at bregma −2 mm AP). The marked region of the skull is then removed using a dental drill. The surgical site is cleaned with saline and sterilized coverslip is placed on the dural surface and a layer of cyano-acrylic glue is put across the perimeter of the cover slip such that a window of 3mm diameter is exposed. An aluminum head bar with one threaded hole is placed on the anterior part of the exposed skull and fixed using an adhesive. Next, dental cement is applied over the exposed skull using sterile toothpick. The mice are then maintained on antibiotics and daily injections of carprofen and dexamethasone for 1 week. Mice recovered for at least 1 week before habituation.

### Two Photon Imaging

A custom-built two-photon laser scanning microscope was built with a Spectra-Physics Tsunami femtosecond laser. Femtosecond pulses centered at 910 nm is coupled to an upright microscope (Zeiss Axio-Examiner.A1) after passing through galvonometric scanner. A 25X 1.05 NA multiphoton excitation optimized water immersion objective (Olympus XLPLN25XWMP2) is used to acquire images. PMT modules from Hamamatsu Corporation (H7422) is used to detect fluorescence. Mice are anesthetized with isoflurane in the holding chamber, attached to a head mount and then transferred to the custom translational head-stage below the objective. In the first imaging session, vein images are obtained using the widefield imaging port and the head-stage co-ordinates are recorded to aid longitudinal imaging of the same regions. 4 Rois of size ~200μm*200μm*150μm and 1024*1024*150 pixels were imaged from each mouse using *ScanImage (r3.8.1)* software. The same ROIs are imaged repeatedly across experimental days. The laser power used was less than 20mW to avoid photodamage.

### Image and Data analysis

Dendritic spines are analyzed using ImageJ and Specifically, the *Simple neurite tracer* plugin included in the FiJi package. An entire Roi is opened in the simple neurite tracer and individual dendrites and spines are traced. Individual dendritic segments with spines are skeletonized and saved as MIP projection images. The tracing and skeletonization of dendritic segments are done by person blind to the training status. A subset of the images is traced by two experimenters independently to confirm the results. The MIP projections are used to quantify presence, gain and loss of spines using custom Matlab code. The distances between the spines are measured by creating geodesic map of the MIP image using matlab inbuilt function *bwdistgeodesic*. The same geodesic map is used to create the digital sequence of dendrites with 1’s representing presence of spines and number of units represents the true length of the dendritic segments. The autocorrelation function of the digital sequence was generated using Matlab command *xcorr*. The ACF data is then binned using a custom code written in JAVA that implements the log binning^2^.

### Statistics

All statistical analysis was done using Origin version 2020b.Behaviour was carried out on 9 adult male mice for Group 1 and 8 mice for group 2 (Fig 1A). Freezing data collected on these groups are analysed using ANOVA and post hoc Bonferoni test for statistical differences. Turnover of the spine data between imaging sessions within a group (3 mice per group, Fig 1B) was analysed using ANOVA and post doc Bonferoni test. Model free empirical differences in ACF values collected per imaging session (ACF of ~300 dendrites) was done using non-parametric Friedman ANOVA. ACF values of data collected from each imaging sessions is compared to ACFs of shuffled data using Friedman ANOVA. Post hoc analysis of ACF values were carried out using Dunn’s test (Ref: Table 1). Non-linear least squares fitting was carried out in Origin through built-in Levenberg Marquet algorithms for minimizing the chi-sq. All the fits with an Adjusted R-Square < 0.7 were considered to not fit the model used and were not used for parameter comparison. Statistical comparison of the fit parameters was carried out using AIC and F-test (Ref: Table 2–5) to establish the differences using Origin. The comparison algorithm is built-in in Origin (https://www.originlab.com/fileExchange/details.aspx?fid=302) briefly we are reproducing the logic and the definitions below:

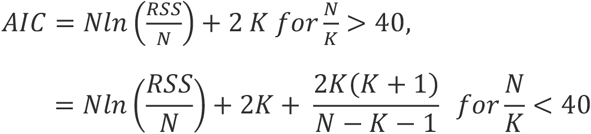

AIC Weight:

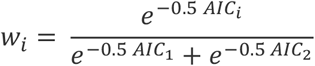

F-Test:

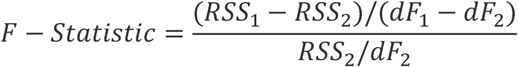

RSS – Residual Sum Square, N – number of data points, K - number of free/fit parameters, dF – degree of freedom. When comparing parameters across two fits, the parameters are individually compared. For comparison the data sets (two of them at any given time) are fit to a given function following two procedures a) two independentl fits with their own free parameter and b) global fit with the parameter in comparison being shared. The RSS obtained is then used to perform AIC and F-Tests. Higher AIC weight (>0.5) indicates the loss in information is lesser for independent fits and parameters are different across the fits. In F-Test p value < 0.05 is taken to be indicative of difference between the parameters being significant. The general logic behind both these methods of comparison being that if the global fit is better (as measured through test statistic (AIC or F-statistic)) then the difference in the value of the fit parameter we obtain is superfluous. The datasets do not need two different values of the parameter that is being compared. The AIC weight quantitatively measures the degree of the information loss that is reduced by having different values for the parameter in question while F-Test yields the measure of the parameters being different through maximum likelihood.

---

**SFigure 1:**
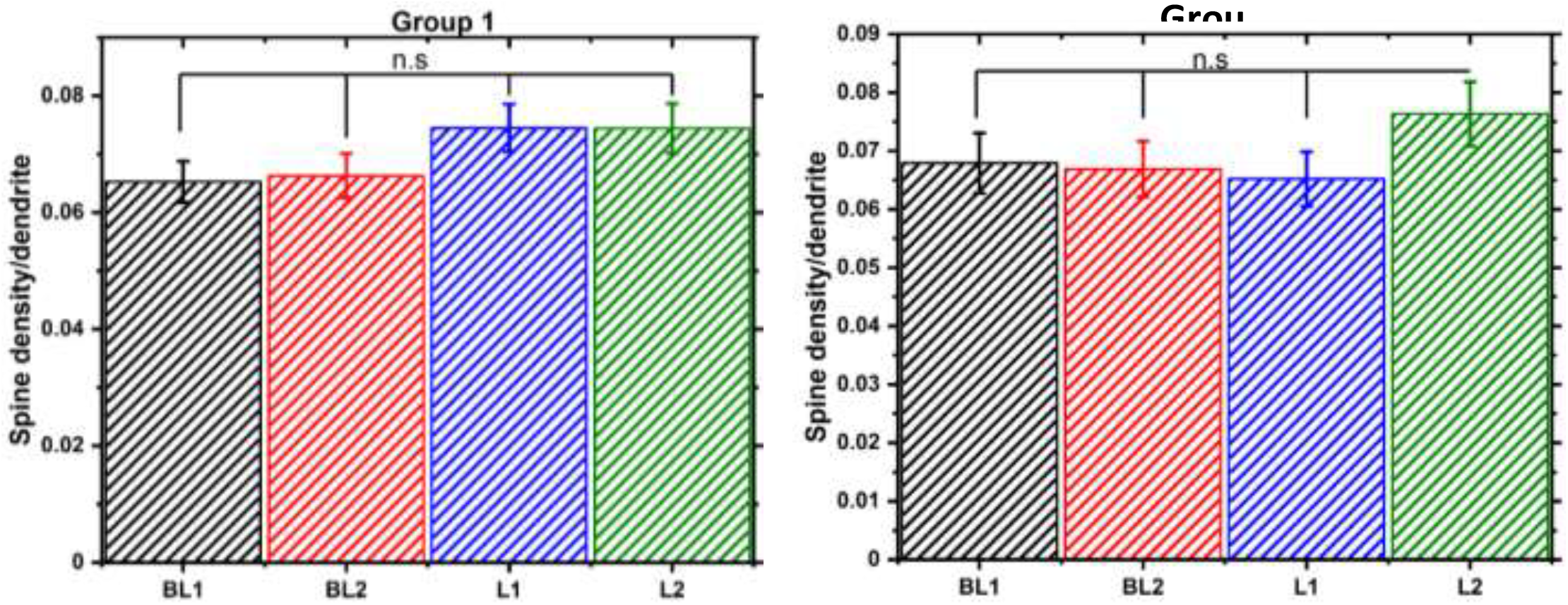
Spine density of Grp1 and Grp2 animals measured during sessions BL1, BL2, L1 and L2. The densities are compared using ANOVA and we find the differences to be non-significant.

**SFigure 2:**
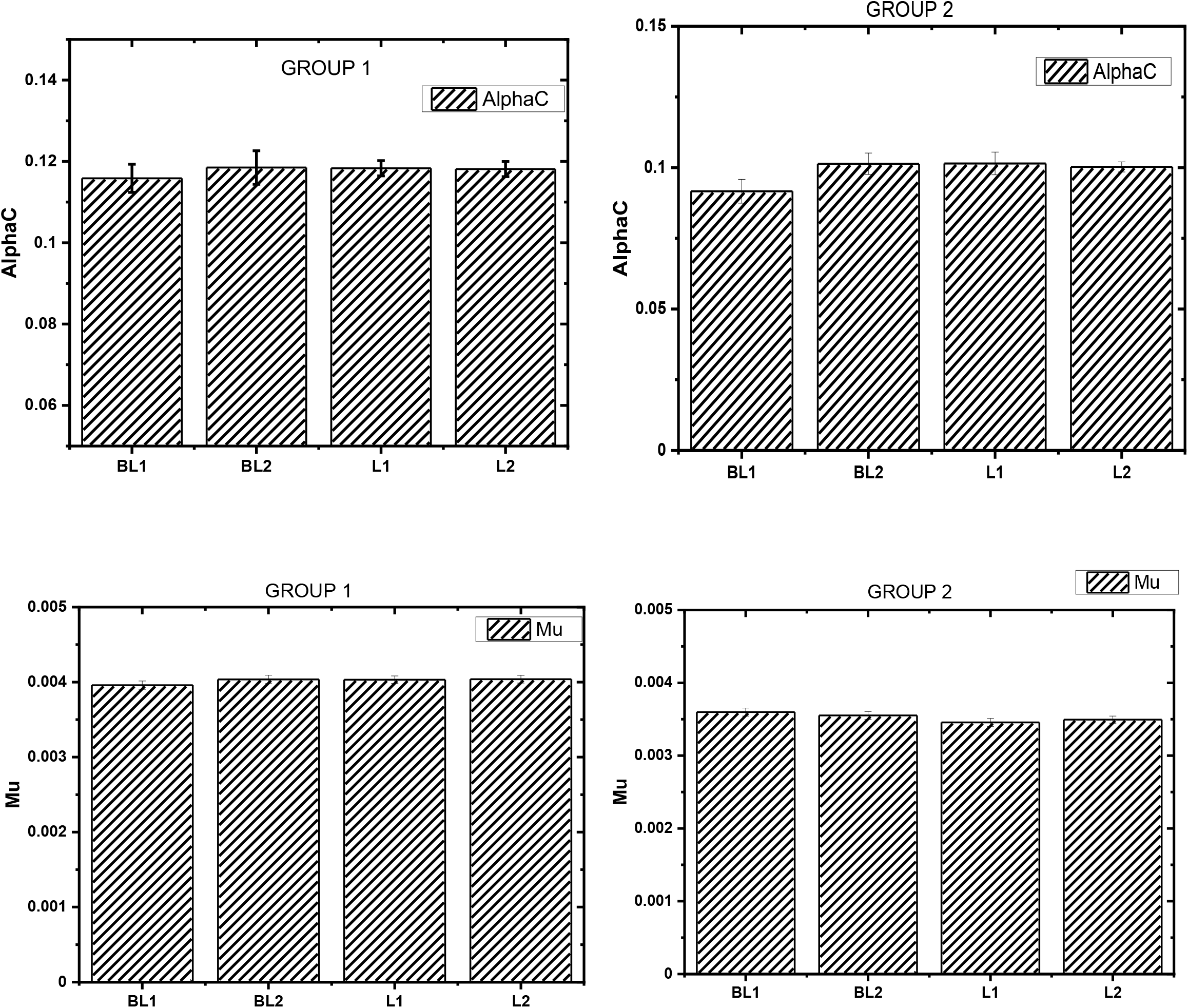
Fit parameters of Grp1 AND Grp2 animals when the unscaled data is fit to Eq.3.

**SFigure 3:**
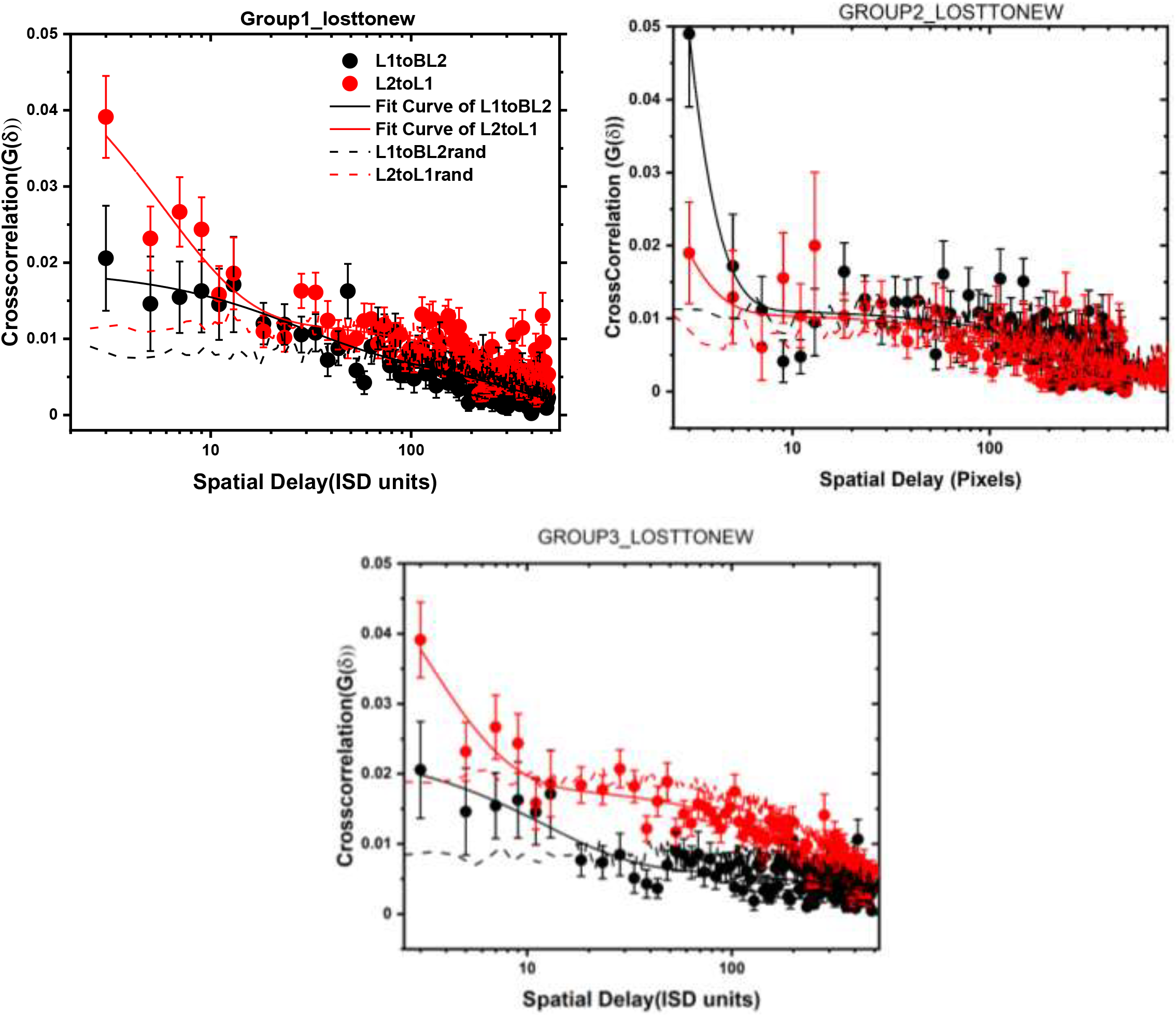
Cross correlation of lost and new dendritic sequence compared with baseline and shuffled.

**SFigure 4:**
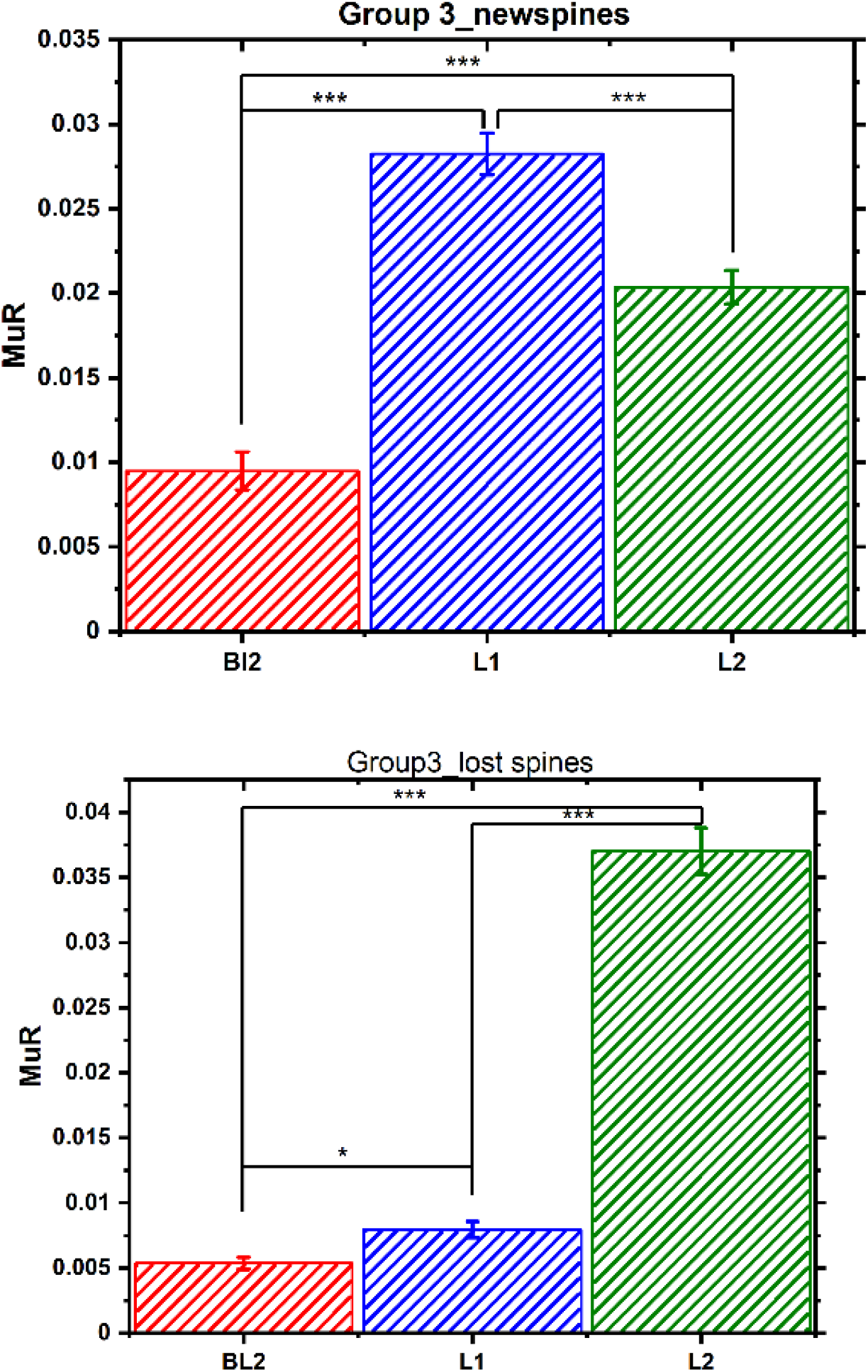
Relative Spine Density of Independent Fraction that exhibits random correlation in new and lost spines of Group 3 animals. The fit parameter μ_R_ obtained from scaled ACFs in each session is shown as bar graphs.

**SFigure 5:**
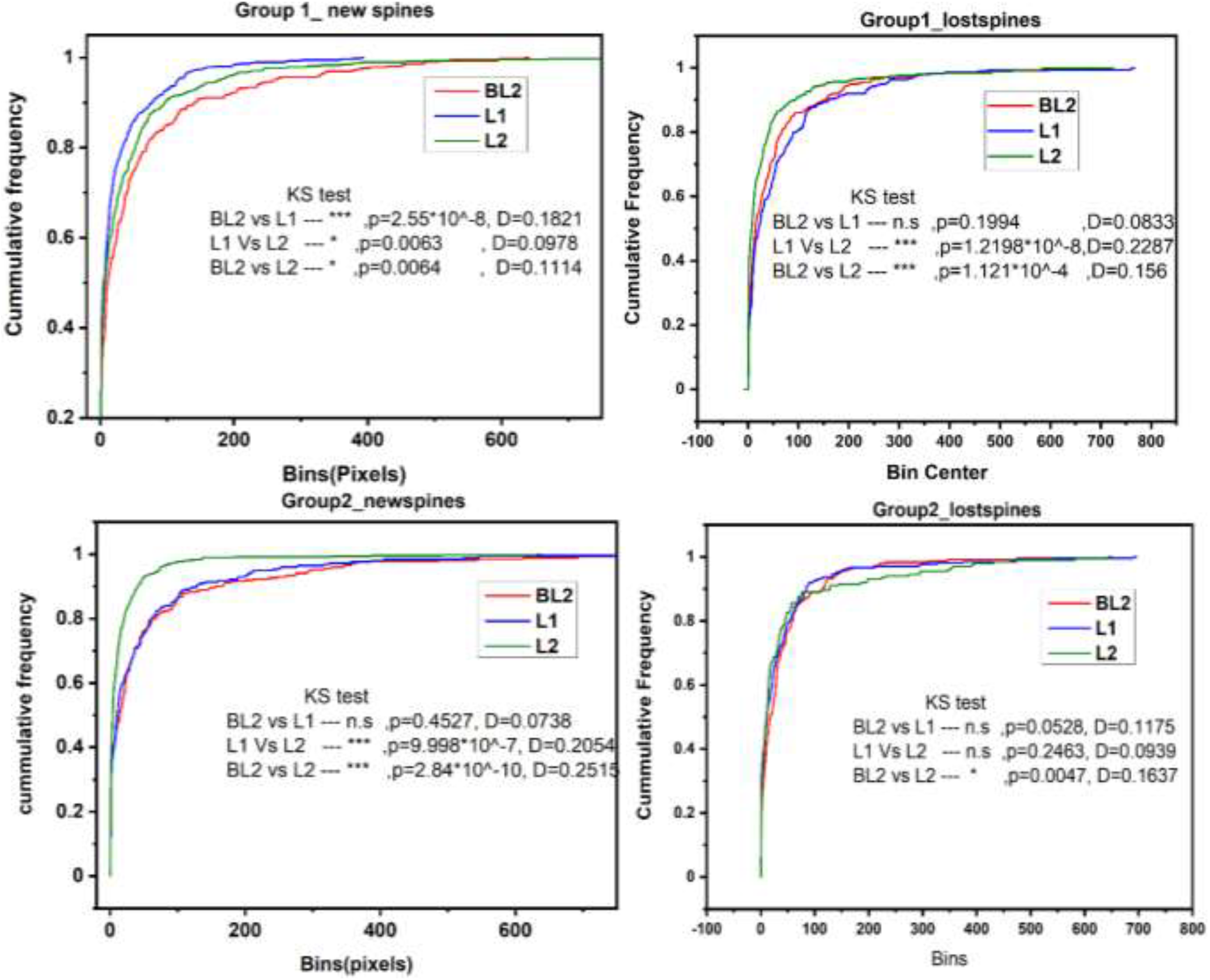
Cumulative Density Function of distance between new spines added(left panel), spines that are lost (right panel) in Group1 (top panel) and Group 2 (bottom panel) animals across imaging sessions. The redlines are the CDFs measured following second baseline (new/lost spines are identified by comparing baseline 1), blues lines are the CDF obtained following L1 (comparing with BL2) and green lines are the CDFs obtained following L2.

**SFigure 6:**
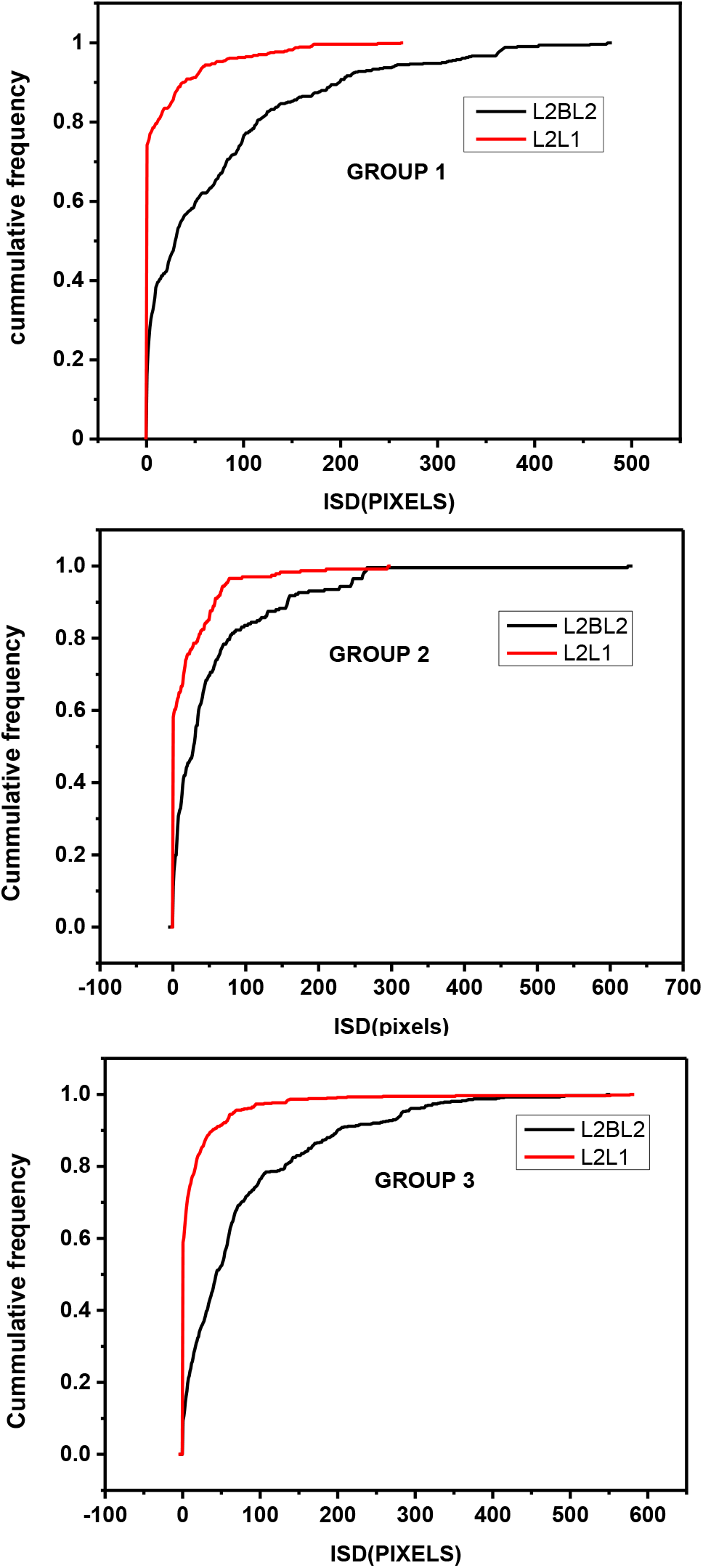
Cumulative Density Function of distance between location of spine loss and next neighbouring spine that was added in the previous session. Group1(Top left), Group 2(Top right) and Group 3(bottom left) animals are shown. The redlines are the CDFs measured by comparing lost spine in second learning session with (i) new spines formed during first learning or (ii) new spines formed during second baseline session (black solid line).

## Notes

### Competing Interest Statement

The authors have declared no competing interest.

